# Omicron-B.1.1.529 leads to widespread escape from neutralizing antibody responses

**DOI:** 10.1101/2021.12.03.471045

**Authors:** Wanwisa Dejnirattisai, Jiandong Huo, Daming Zhou, Jiří Zahradník, Piyada Supasa, Chang Liu, Helen M.E. Duyvesteyn, Helen M. Ginn, Alexander J. Mentzer, Aekkachai Tuekprakhon, Rungtiwa Nutalai, Beibei Wang, Aiste Dijokaite, Suman Khan, Ori Avinoam, Mohammad Bahar, Donal Skelly, Sandra Adele, Sile Ann Johnson, Ali Amini, Thomas Ritter, Chris Mason, Christina Dold, Daniel Pan, Sara Assadi, Adam Bellass, Nikki Omo-Dare, David Koeckerling, Amy Flaxman, Daniel Jenkin, Parvinder K Aley, Merryn Voysey, Sue Ann Costa Clemens, Felipe Gomes Naveca, Valdinete Nascimento, Fernanda Nascimento, Cristiano Fernandes da Costa, Paola Cristina Resende, Alex Pauvolid-Correa, Marilda M. Siqueira, Vicky Baillie, Natali Serafin, Zanele Ditse, Kelly Da Silva, Shabir Madhi, Marta C Nunes, Tariq Malik, Peter JM Openshaw, J Kenneth Baillie, Malcolm G Semple, Alain R Townsend, Kuan-Ying A. Huang, Tiong Kit Tan, Miles W. Carroll, Paul Klenerman, Eleanor Barnes, Susanna J. Dunachie, Bede Constantinides, Hermione Webster, Derrick Crook, Andrew J Pollard, Teresa Lambe, OPTIC consortium, ISARIC4C consortium, Neil G. Paterson, Mark A. Williams, David R. Hall, Elizabeth E. Fry, Juthathip Mongkolsapaya, Jingshan Ren, Gideon Schreiber, David I. Stuart, Gavin R Screaton

**Affiliations:** Wellcome Centre for Human Genetics, Nuffield Department of Medicine, University of Oxford, Oxford, UK; Division of Structural Biology, Nuffield Department of Medicine, University of Oxford, The Wellcome Centre for Human Genetics, Oxford, UK; Chinese Academy of Medical Science (CAMS) Oxford Institute (COI), University of Oxford, Oxford, UK; Department of Biomolecular Sciences, Weizmann Institute of Science, Rehovot, Israel; Diamond Light Source Ltd, Harwell Science & Innovation Campus, Didcot, UK; Oxford University Hospitals NHS Foundation Trust, Oxford, UK; Peter Medawar Building for Pathogen Research, Oxford, UK; Nuffield Department of Clinical Neurosciences, University of Oxford, Oxford, UK; Translational Gastroenterology Unit, University of Oxford, Oxford, UK; NIHR Oxford Biomedical Research Centre, Oxford, UK; Oxford Vaccine Group, Department of Paediatrics, University of Oxford, Oxford, UK; Department of Infectious Diseases and HIV Medicine, University Hospitals of Leicester NHS Trust; Department of Respiratory Sciences, University of Leicester; Medical Sciences Division, University of Oxford; Jenner Institute, Nuffield Department of Medicine, University of Oxford, Oxford, UK; Institute of Global Health, University of Siena, Siena, Brazil; Department of Paediatrics, University of Oxford, Oxford, UK; Laboratório de Ecologia de Doenças Transmissíveis na Amazônia, Instituto Leônidas e Maria Deane, Fiocruz, Manaus, Amazonas, Brazil; Fundação de Vigilância em Saúde do Amazonas, Manaus, Amazonas, Brazil; Laboratorio de vírus respiratórios- IOC/FIOCRUZ, Rio de Janeiro, Brazil; Department of Veterinary Integrative Biosciences, Texas A&M University, College Station, TX, United States; South African Medical Research Council, Vaccines and Infectious Diseases Analytics Research Unit, School of Pathology, Faculty of Health Sciences, University of the Witwatersrand, Johannesburg, South Africa; Department of Science and Technology/National Research Foundation, South African Research Chair Initiative in Vaccine Preventable Diseases, Faculty of Health Sciences, University of the Witwatersrand, Johannesburg, South Africa; National Infection Service, Public Health England (PHE), Porton Down, Salisbury, UK; National Heart & Lung Institute, Imperial College London; Genetics and Genomics, Roslin Institute, University of Edinburgh, Edinburgh, UK; NIHR Health Protection Research Unit, Institute of Infection, Veterinary and Ecological Sciences, Faculty of Health and Life Sciences, University of Liverpool, Liverpool, UK; MRC Human Immunology Unit, Weatherall Institute of Molecular Medicine, University of Oxford, John Radcliffe Hospital, Oxford, OX3 9DS, UK; Research Center for Emerging Viral Infections, College of Medicine, Chang Gung University, Taoyuan, Taiwan; Centre For Tropical Medicine and Global Health, Nuffield Department of Medicine, University of Oxford, Oxford, UK; Mahidol-Oxford Tropical Medicine Research Unit, Bangkok, Thailand, Department of Medicine, University of Oxford, Oxford, UK; Nuffield Department of Medicine, University of Oxford, Oxford, UK; Siriraj Center of Research Excellence in Dengue & Emerging Pathogens, Dean Office for Research, Faculty of Medicine Siriraj Hospital, Mahidol University, Thailand; Instruct-ERIC, Oxford House, Parkway Court, John Smith Drive, Oxford, UK

## Abstract

On the 24^th^ November 2021 the sequence of a new SARS CoV-2 viral isolate spreading rapidly in Southern Africa was announced, containing far more mutations in Spike (S) than previously reported variants. Neutralization titres of Omicron by sera from vaccinees and convalescent subjects infected with early pandemic as well as Alpha, Beta, Gamma, Delta are substantially reduced or fail to neutralize. Titres against Omicron are boosted by third vaccine doses and are high in cases both vaccinated and infected by Delta. Mutations in Omicron knock out or substantially reduce neutralization by most of a large panel of potent monoclonal antibodies and antibodies under commercial development. Omicron S has structural changes from earlier viruses, combining mutations conferring tight binding to ACE2 to unleash evolution driven by immune escape, leading to a large number of mutations in the ACE2 binding site which rebalance receptor affinity to that of early pandemic viruses.

## Introduction

Since the end of 2020 a series of viral variants have emerged in different regions where some have caused large outbreaks, Alpha (Supasa et al., 2021) and more recently Delta (Liu et al., 2021a) have had the greatest global reach, whilst Beta (Zhou et al., 2021), Gamma (Dejnirattisai et al., 2021b) and Lambda (Colmenares-Mejia et al., 2021), although causing large outbreaks in Southern Africa and South America, did not become dominant in other parts of the World. Indeed, Beta was later displaced by Delta in South Africa.

The rapid emergence of Omicron (https://www.who.int/news/item/26-11-2021-classification-of-omicron-(bi1.1.529)-sars-cov-2-variant-of-concern) on the background of high Beta immunity implies that the virus may have evolved to escape neutralization by Beta specific serum (Liu et al., 2021b). Within S, Omicron has 30 substitutions plus the deletion of 6 and insertion of 3 residues, whilst in all the other proteins there are a total of 16 substitutions and 7 residue deletions. Particular hotspots for the mutations are the ACE2 receptor binding domain (RBD) (15 amino acid substitutions) and the N-terminal domain (NTD) (3 deletions totalling 6 residues, 1 insertion, 4 substitutions). In-depth studies by a large number of laboratories have shown the RBD and NTD as the site of binding of the most potent monoclonal antibodies (mAbs) (Cerutti et al., 2021; Dejnirattisai et al., 2021a; Rapp et al., 2021; Zost et al., 2020) and the RBD is the site of binding of various mAbs in clinical development (Baum et al., 2020; Dong et al., 2021; Pinto et al., 2020; Starr et al., 2021).

S mediates cellular interactions. It is a dynamic, trimeric structure (Walls et al., 2020; Walls et al., 2017; Wrapp et al., 2020) which can be lipid bound (Toelzer et al., 2020) and tightly associated in a ‘closed’ form or unfurled to expose one or more RBDs allowing both receptor binding and increased access to neutralising antibodies. Once bound to a cell, S undergoes cleavage and a drastic elongation converting it to the post fusion form.

Most potent neutralizing antibodies target the ACE2 footprint (Dejnirattisai et al., 2021a; Lan et al., 2020; Liu et al., 2021b) occupying ∼880 A^2^ at the outermost tip of the RBD (the neck and shoulders referring to the torso analogy (Dejnirattisai et al., 2021a)), preventing cell attachment. A proportion of antibodies are able to cross-neutralize different variants (Liu et al., 2021b) and a few of these bind to a motif surrounding the N-linked glycan at residue 343 (Dejnirattisai et al., 2021a; Liu et al., 2021b). These antibodies, exemplified by S309 (Pinto et al., 2020) can cross-react with SARS-CoV-1 but do not block ACE2 interaction and their mechanism of action may be to destabilize the S-trimer. Neutralizing anti-NTD mAbs do not block ACE2 interaction and bind to a so-called supersite on the NTD (Cerutti et al., 2021; Chi et al., 2020), however they generally fail to give broad protection as the supersite is disrupted by a variety of NTD mutations present in the variants of concern. Moreover, some NTD binding antibodies were shown to have an infectivity-enhancing effect by induction of S open state (Liu et al., 2021c).

A relation between higher binding affinity of the RBD to ACE2 and higher infectivity has been previously suggested (Starr et al., 2020; Zahradnik et al., 2021b). Indeed, mutations in the RBD of Alpha, Beta, Gamma and Delta variants are associated with an increased affinity towards ACE2 (Dejnirattisai et al., 2021a; Liu et al., 2021a; Supasa et al., 2021; Zahradnik et al., 2021b; Zhou et al., 2021). Interestingly, yeast surface displayed selection of randomly mutated RBD selected for tighter ACE2 binding selected the mutations present in the variants of concern N501Y (Alpha, Beta, Gamma) and E484K (Beta, Gamma), variants of interest F490S (Lambda) or multiple other variants, for instance S477N (Zahradnik et al., 2021b). In addition, the mutation Q498R was selected, and shown to have a major contribution to binding (in epistasis with N501Y), raising the concern that a new variant that includes the different affinity enhancing mutations may arise (Zahradnik et al., 2021b).

In this report, we study the neutralization of Omicron by a large panel of sera collected from convalescent early pandemic, Alpha, Beta, Gamma and Delta infected individuals, together with vaccinees receiving three doses of the Oxford/AstraZeneca (AZD1222) or the Pfizer BioNtech (BNT16b2) vaccines. There is widespread reduction in the neutralization activity of serum from multiple sources and we use these data to plot an antigenic map where Omicron is seen to occupy the most distant position from early pandemic viruses which form the basis for current vaccines.

We show that Omicron escapes neutralization by the majority of potent monoclonal antibodies (mAbs) arising after both early pandemic and Beta variant. Utilizing a large bank of structures (n=29) from panels of potent monoclonal antibodies we describe the mechanism of escape caused by the numerous mutations present in Omicron RBD, which includes most mAbs developed for prophylactic or therapeutic use (Baum et al., 2020).

Analysis of the binding of ACE2 to RBD and structural analysis of the Omicron RBD indicates that changes at residues 498 and 501 of the RBD have locked ACE2 binding to the RBD in that region sufficiently strongly to enable the generation of a plethora of less favourable changes elsewhere, providing extensive immune escape and in the process resulting in a final net affinity for ACE2 similar to the early pandemic virus.

## Results

### Phylogeny of Omicron

On average, a mutation is inserted into the genome of SARS-CoV-2 every second infection. Such mutations generate intra-host diversity upon which selection then acts, with the viral load being a critical parameter (Valesano et al., 2021). As a result, the virus is constantly changing. The median number of sequences that harbor a mutation at any given position is 570 (out of 4.7 million genomic sequences in GISAID; Nov 14, 2021), however rather few of these are enriched, with less than 10% of amino-acids having over 10,000 sequenced mutations. Enrichment of a mutation at a given position suggests increased virus fitness endowing a selective advantage (Martin et al., 2021; Zahradnik et al., 2021c). If a specific mutation arises independently in different lineages, it implies an even greater advantage in fitness.

Omicron has changes throughout its proteome but S changes dominate, with 30 amino acid substitutions plus 6 residues deleted and 3 inserted (**Figure 1 and 2**). Ten of these were found previously in at least two lineages (D614G was mutated early on and maintained throughout). Of those ten, six have the same amino-acid substitution in >75% of the sequences, and only one (E484A) has a unique substitution in Omicron (in Beta and Gamma it is a Lys). **Figure S1A** shows the number of mutant sequences per residue at positions undergoing mutations in independent lineages. This can be interpreted in two ways, one is that the later mutations are epistatic to one another and thus are more difficult to reach, or that they do not contribute to virus fitness.

**Figure 1.**
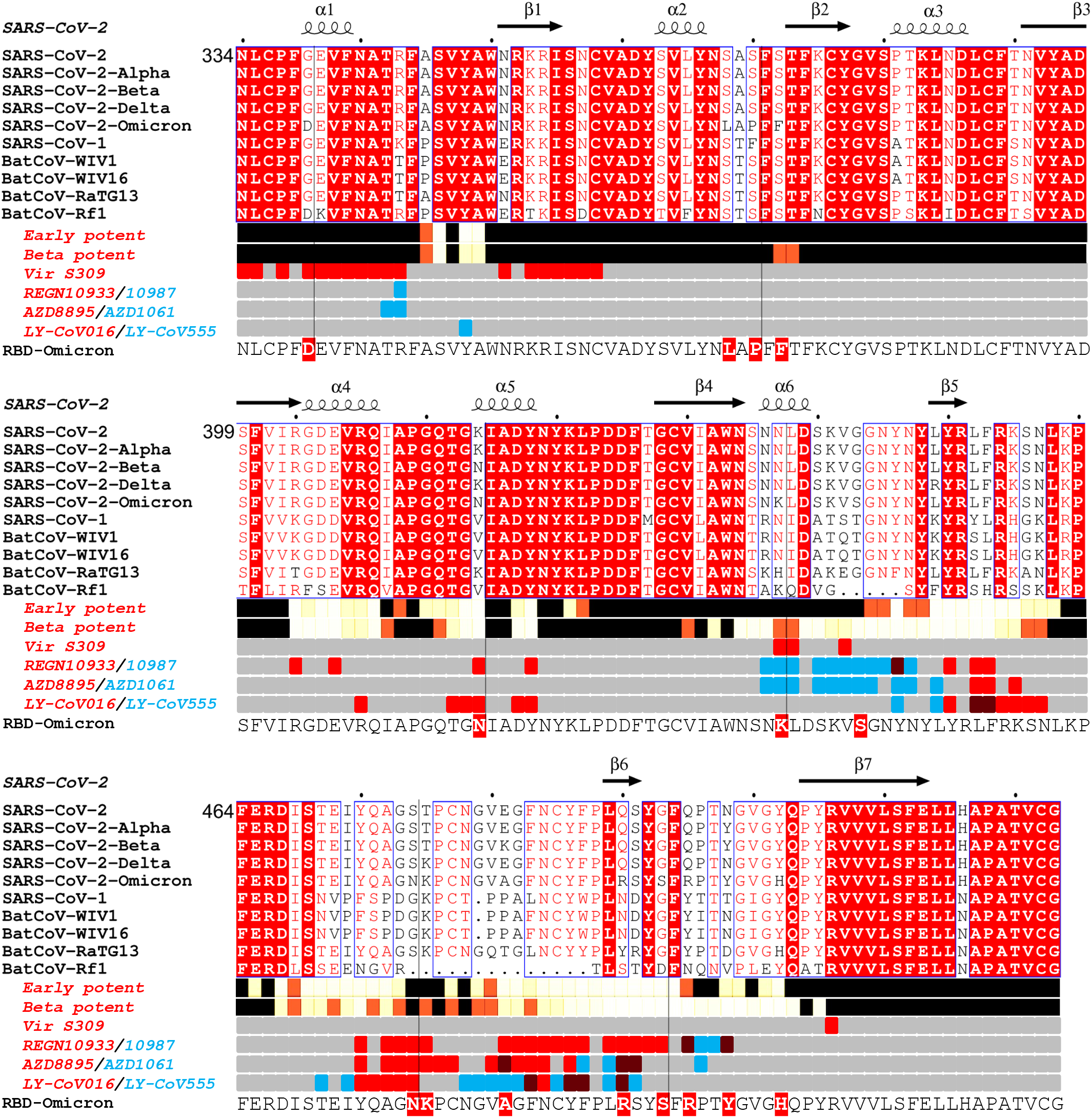
Sarbecovirus RBD sequence analysis. Shown with Alpha, Beta, Delta and Omicron variants (the latter repeated on the lower line to clarify the Omicron changes. Binding sites for the early pandemic potent antibodies (Dejnirattisai et al., 2021a) and the potent Beta antibodies ((Liu et al., 2021b) are depicted using iron heat colours (grey > straw > blue > glowing red > yellow > white) to indicate relative levels of antibody contact and commercial antibody contacts are depicted with the pairs of antibodies in red or blue with purple denoting interactions with the same residue). Totally conserved residues are boxed on a red background on the upper rows, whilst on the final row the Omicron mutations are boxed in red. Secondary elements are denoted above the alignment. The figure was produced in part using Espript (Robert and Gouet, 2014).

**Figure 2.**
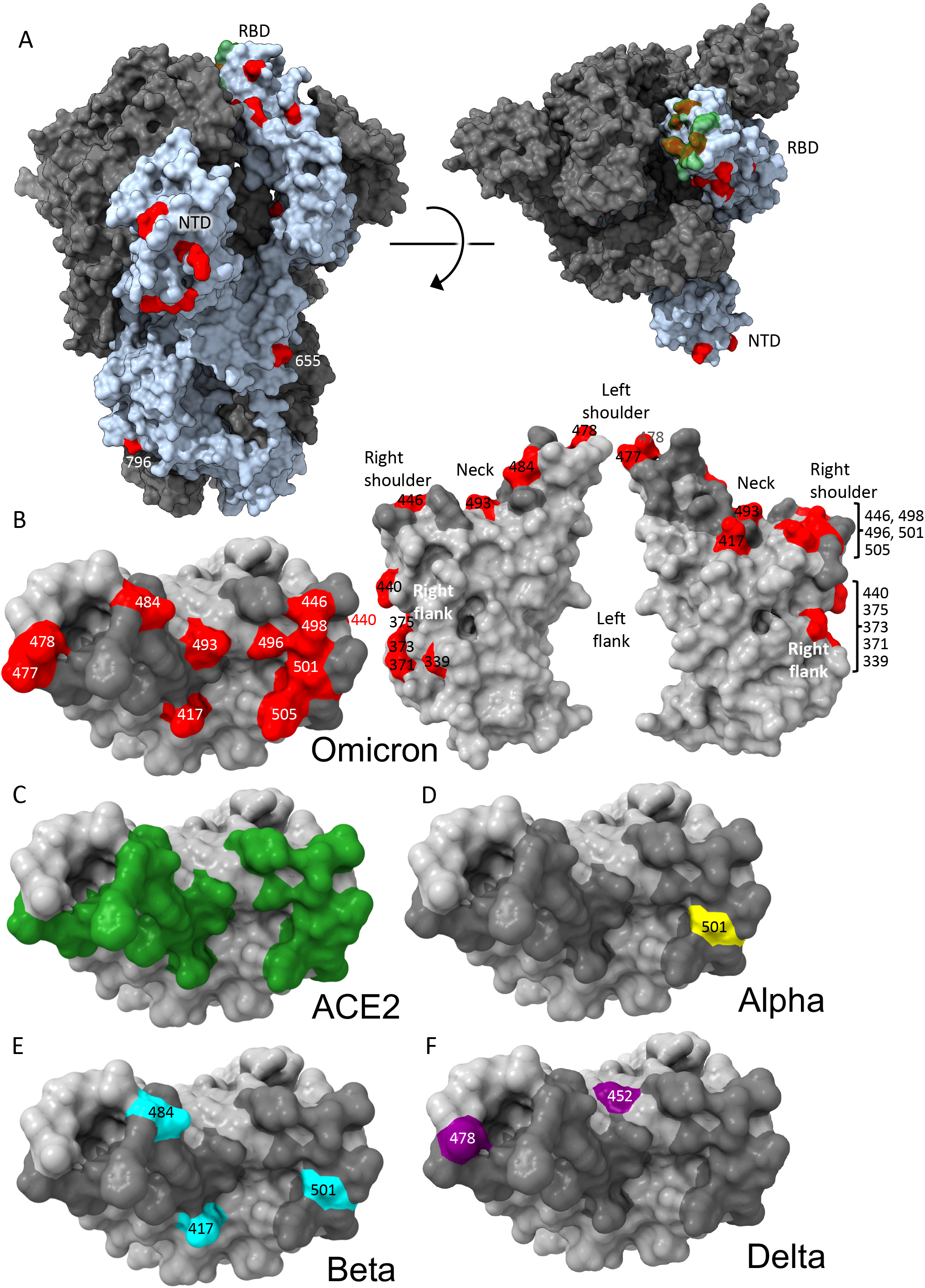
Distribution of Omicron changes. (A) Trimeric S model depicted as a grey surface with one monomer highlighted in pale blue, ACE2 binding site in green and changes in Omicron shown in red, left side view, right top view. (B) RBD depicted as a grey surface with the ACE2 footprint in dark grey and changes in Omicron in red, left: top view, right: front and back views. Epitopes are labelled according to the torso analogy and mutations labelled. (C,D,E,F) Top view of RBD depicted as a grey surface with (C), ACE2 binding site in green (D), Alpha change in yellow (E), Beta changes in cyan (F), Delta changes in purple. Figure produced using chimera X (Pettersen et al., 2021).

The Omicron RBD has 15 changes in total in the RBD, described below. The NTD also has numerous changes, including 4 amino acid substitutions, 6 amino acids deleted and 3 amino acids inserted, also described below. Several mutations found in Omicron occur in residues conserved in SARS-CoV-1 and many other Sarbecoviruses (**Figure 1**). These observations agree with the Pango classification (Rambaut et al., 2020) which places Omicron at a substantial distance from all other variants.

Although S has the largest concentration of changes, there are mutations in many non-structural proteins. Nsp12 Polymerase (PDB:7B3B) (Gao et al., 2020) bears a single mutation in Omicron, P323L, distant from the active site targeted by compounds in use or in clinical trials, whilst the Nsp14 N-terminal exonuclease domain (PDB:7N0D) (Moeller et al., 2021) important in proofreading, again has a single mutation, I42V, distal to the active site. These observations suggest that the large number of mutations in Omicron is not being driven by reduced polymerase fidelity. Nsp5 Mpro protease (PDB:7RFW) (Owen et al., 2021) has a single mutation, P132H, distal to the active site targeted by the Pfizer PF-07321332 compound and Nsp3 has no mutations close to either the active site of the papain-like protease (Gao et al., 2021) or the macro domain inhibitor site. Whilst allosteric effects cannot be completely ruled out, there is little apparent significance to these changes which probably would not impact on small molecule therapeutics currently in use or development. This may reflect the lack of selective pressure to date on these proteins.

### Mapping of Omicron RBD mutations compared to Alpha, Beta, Gamma and Delta

The Alpha variant has a single change in the RBD at N501Y (**Figure 2D**) (Supasa et al., 2021) which occupies the right shoulder and contributes to the ACE2 binding footprint. Beta has two further mutations in the RBD: K417N and E484K, at back of the neck and left shoulder respectively (**Figure 2E**), also part of the ACE2 footprint (**Figure 2C**) (Zhou et al., 2021). Gamma mutations are similar: K417T, E484K, N501Y (Dejnirattisai et al., 2021b). Delta mutations, L452R front of neck, T478K far side of left shoulder, fall just peripheral to the ACE2 binding footprint (**Figure 2F**) (Liu et al., 2021a). All of these variants have at least one RBD mutation in common with Omicron. Of the 15 Omicron changes in the RBD, nine map to the ACE2 binding footprint: K417N, G446S, S477N, E484A, Q493R, G496S, Q498R, N501Y, Y505H with N440K and T478K just peripheral (**Figure 2B-C**). Aside from these, mutations occur on the right flank: G339D, S371L, S373P and S375F (**Figure 2B**), the last three of which are adjacent to a lipid binding pocket (**Figure S1B**) (Toelzer et al., 2020). The lipid binding pocket has been seen occupied by a lipid similar to linoleic acid in an unusually rigid state of S where all RBDs are found in a locked-down configuration stabilised by lipid-bridged quaternary interactions between adjacent RBDs. However, this lipid bound form has been rarely seen, instead the pocket is usually empty and collapsed, with the RBD alternating between looser down and up conformations. We presume that this is because the pocket readily empties of lipid during protein purification, indeed rapidly prepared virus particles tend to have the RBDs closer to the locked down state (Ke et al., 2020). Loss of lipid promotes RBD presentation to the target cell.

Until now the antigenic properties of variant viruses have been well described by assuming each change produces only a local change in structure and we will use this assumption in rationalising the serological impact of the changes in Omicron. We present structural data later to qualify this assumption.

### Omicron NTD mutations

The heavily mutated Omicron NTD possesses the 3 deletions found in Alpha, Delta has 1/3 substitutions in common with Omicron, whilst there are no NTD changes in common with Beta or Gamma. The mutations seen in the NTD lie on exposed flexible loops, which differ from those in SARS-CoV-1 and are likely to be antigenic (**Figure 1A**). In summary, the pattern of deletions and insertions seen in Omicron consistently moves those loops that are most different from SARS-CoV-1 to being more SARS-CoV-1-like, at least in length. Of the N1, N3 and N5 loops which comprise the antibody supersite, Omicron has a substitution at G142D and deletion of residues 143-145 in N3 which would mitigate against binding by a number of potent neutralizing antibodies e.g. 4A8 and 159 (Chi et al., 2020; Dejnirattisai et al., 2021b). The deletion of residues 69 and 70 in N2 has also occurred in the Alpha variant whilst the deletion at residue 211, substitution at 212 and insertion at 214 are unique to Omicron. All these changes are on the outer surface and likely antigenic.

### Neutralization of Omicron by Convalescent serum

We isolated Omicron virus from the throat swab of an infected case in the UK. Following culture in VeroE6 cells transfected with TMPRSS2 the S gene sequence was confirmed to be the Omicron consensus with the additional mutation A701V, which is present in a small number of Omicron sequences.

We have collected sera from individuals infected early in the pandemic (n=32) before the emergence of the variants of concern (VOC), along with cases infected with Alpha (n=18), Beta (n=14), Gamma (n=16) and Delta (n=42). Neutralization assays were performed against Omicron and compared with neutralization titres for Victoria (an early pandemic strain), Alpha, Beta, Gamma and Delta.

In all cases neutralization titres to Omicron were substantially reduced compared to either the ancestral strain Victoria or to the homologous strain causing infection and in a number of cases immune serum failed to neutralize Omicron at 1/20 dilution (**Figure 3A-E**). Compared to Victoria the neutralization titres of sera for Omicron were reduced for early pandemic 16.9- fold (p<0.0001), Alpha 33.8-fold (p<0.0001), Beta 11.8-fold (p=0.0001), Gamma 3.1-fold (p=0.001) and Delta 1.7-fold (p=0.0182). Compared to the neutralization of homologous virus, for example Alpha virus by Alpha serum, Omicron neutralization was reduced for sera from Alpha 18.4-fold (p<0.0001), Beta 22.5-fold (<0.0001), Gamma 12.3-fold (p<0.0001) and Delta 25.9-fold (p<0.0001).

**Figure 3.**
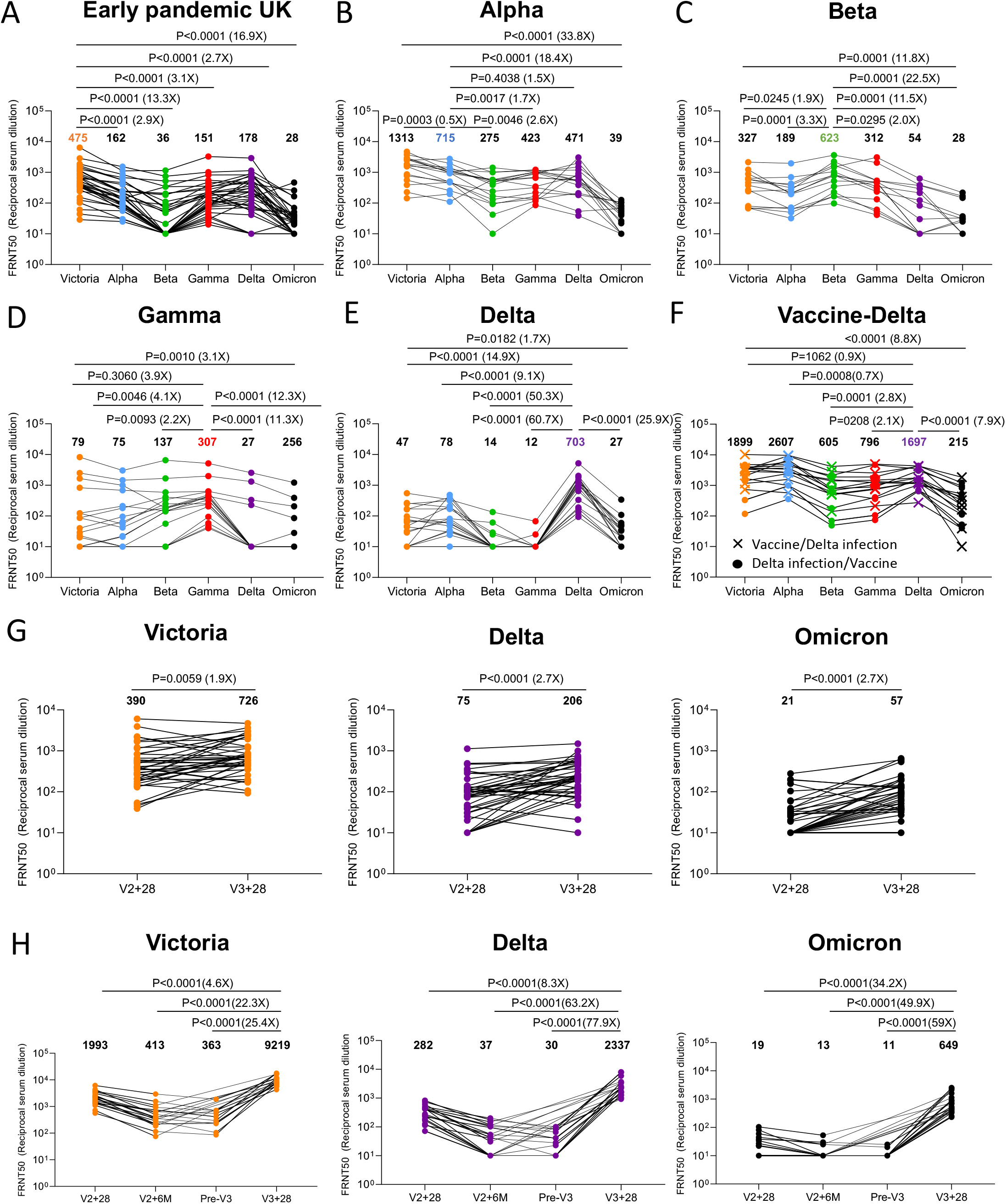
Neutralization assays against Omicron. FRNT50 values for the indicated viruses using serum from convalescent subjects previously infected with A) Early pandemic virus (n=32), (B) Alpha (n=18), (C) Beta (n=14), (D) Gamma (n=16), (E) Delta (n=19), (F) Delta before vaccination or Delta after vaccination (n=17), (G) Before and after the third dose of AZD1222 (n=41), (H) 4 weeks, 6 months after the second dose, before the third and after the third dose of BNT162b2 (n=20). In A-E comparison is made with neutralization titres to Victoria, Alpha, Beta and Gamma and Delta previously reported in (Dejnirattisai et al., 2021a; Supasa et al., 2021; Zhou et al., 2021; Dejnirattisai et al., 2021b; Liu et al., 2021b), in G the data points for BNT162b2 are taken from (Flaxman et al., 2021). Geometric mean titres are shown above each column. The Wilcoxon matched-pairs signed rank test was used for the analysis and two-tailed P values were calculated.

In summary, Omicron causes widespread escape from neutralization by serum obtained following infection by a range of SARS-CoV-2 variants meaning that previously infected individuals will have little protection from infection with Omicron, although it is hoped that they will still maintain protection from severe disease.

### Vaccinated and infected sera show better neutralization of Omicron

We have collected sera from Delta infected cases and because Delta spread in the UK during the vaccination campaign, we obtained sera from three different groups; Delta infection only (n=19) (**Figure 3E**), Delta infection following vaccination (n=9) or vaccination following Delta infection (n=8) (**Figure 3F**). Neutralization assays against early pandemic, Alpha, Beta, Gamma, Delta and Omicron viruses were performed. Compared to Delta-only infected individuals, sera from cases who had received vaccine and been infected by Delta showed substantially higher neutralization to all viruses tested; early pandemic, with Delta infected and vaccinated sera showing a 7.9-fold (p<0.0001) increase in the neutralization of Omicron, compared to Delta infection alone. To confirm the boosting effect of vaccination we collected a paired blood sample from 6 Delta cases before and after vaccination which clearly demonstrates the boosting effect of infection and vaccination (**Figure S2**).

In summary, sera taken from convalescent cases previously infected with a variety of SARS CoV-2 variants show substantial reduction in neutralization titres to Omicron, likely indicating that these individuals will have little protection from reinfection with Omicron, although it is hoped they will retain some protection from severe disease. On the other hand, infection with Delta (and perhaps other variants) together with vaccination significantly boosts Omicron neutralization titres.

### Increased neutralization of Omicron by third dose booster vaccination

In a number of countries such as the UK, booster programmes have been launched to counter waning immunity and the increasing frequency of break-through infections with Delta. To examine the effect of booster vaccination, we tested neutralization of Victoria, Delta and Omicron viruses using sera from individuals receiving 3 doses of ADZ1222 (n=41) or BNT162b2 (n=20). For ADZ1222, serum was obtained 28 days following the second and third doses (**Figure 3G**). For BNT162b2, serum was obtained 28 days, 6 months, immediately prior to the third dose and 28 days following the third dose (**Figure 3H**).

At 28 days following the third dose, for ADZ1222, the neutralization titre to Omicron was reduced 12.7-fold (p<0.0001) compared to Victoria and 3.6-fold (p<0.0001) compared with Delta, for BNT162b2, the neutralization titre to Omicron was reduced 14.2-fold (p<0.0001) compared to Victoria and 3.6-fold (p<0.0001) compared to Delta. The neutralization titres for Omicron were boosted 2.7-fold (p<0.001) and 34.2-fold (P<0.001) following the third dose of ADZ1222 and BNT162b2 respectively compared to 28 days following the second dose. Of concern, and as has been noted previously, neutralization titres fell substantially between 28 days and 6 months following the second dose of the BNT162b2 vaccine, although we did not measure titres 6 months following the second dose of AZD1222.

In summary, neutralization titres against Omicron are boosted following a third vaccine dose, meaning that the campaign to deploy booster vaccines should add considerable protection against Omicron infection.

### Effect of Omicron mutations on Wuhan antibody responses

We have previously reported a panel of 20 potent neutralizing antibodies (FRNT50 < 100ng/ml) isolated from cases infected with early pandemic viruses (Wuhan) (Dejnirattisai et al., 2021a). Neutralization assays against Omicron were performed and compared with neutralization of early pandemic, Alpha, Beta, Gamma, Delta viruses; 17/20 mAbs failed to neutralize Omicron (FRNT50 >10μg/ml) whilst the titres for mAbs 58, 222 and 253 were reduced 3.4, 12.6 and 19.3 -fold compared to Victoria (**Figure 4, Table S1**).

**Figure 4.**
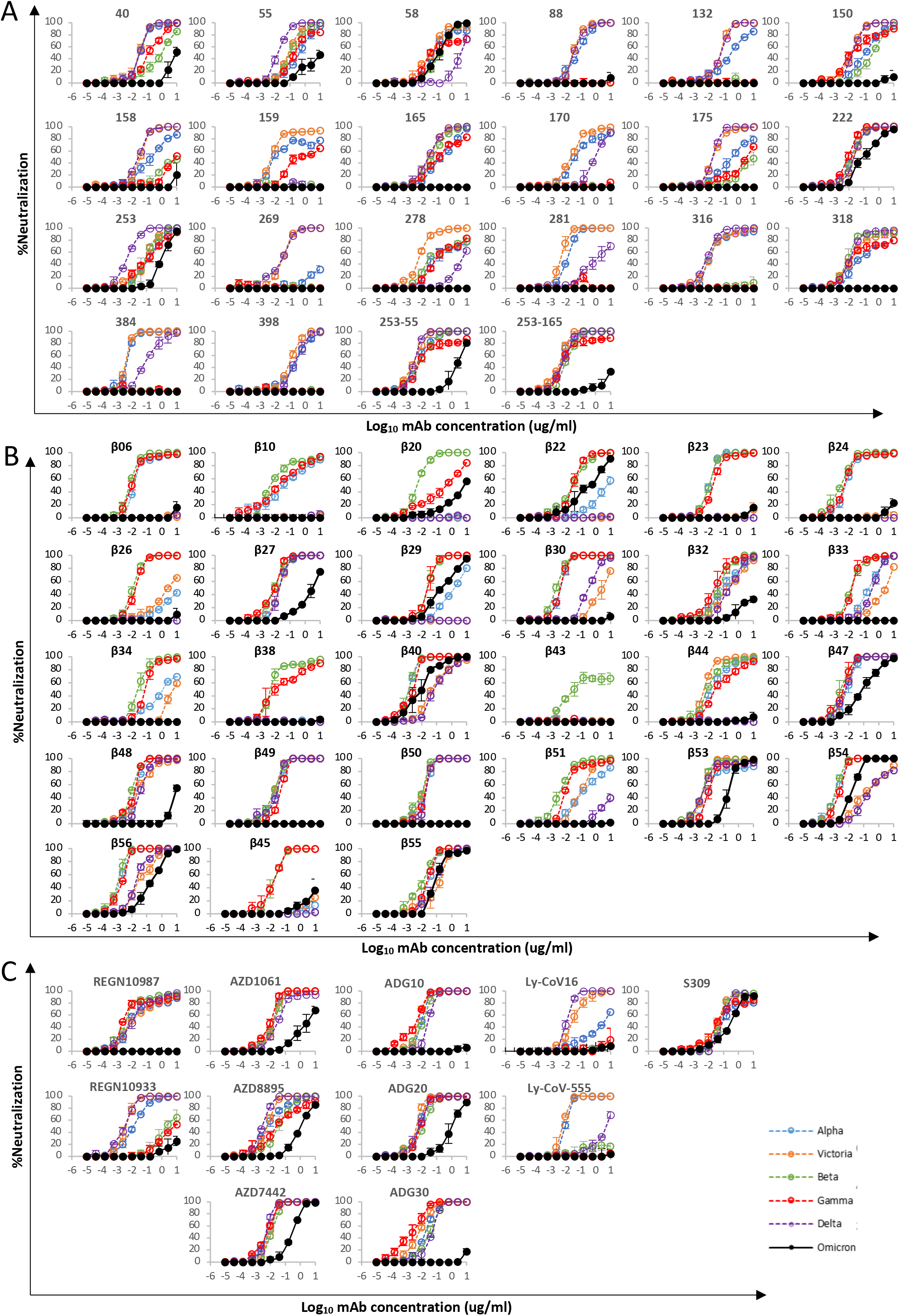
mAb Neutralization curves. FRNT curves for mAb from (A) Early pandemic, (B) Beta infected cases or (C) Commercial sources. Omicron neutralization is compared with curves for Victoria, Alpha, Beta, Gamma and Delta which have been previously reported (Dejnirattisai et al., 2021a; Supasa et al., 2021; Zhou et al., 2021; Dejnirattisai et al., 2021b; Liu et al., 2021b). Neutralization titres are reported in Table S1.

The binding sites of these antibodies were mapped together with other published structures to 5 epitopes (based on the position of the centre of gravity of each antibody) either by direct structural studies or competition analyses (Dejnirattisai et al., 2021a). According to the torso analogy (Dejnirattisai et al., 2021a) these were designated: neck left shoulder, right shoulder, right flank and left flank (**Figure 2B**). In **Figure 5A-D** we show the mapping of the density of centroids to the surface of the RBD with the Omicron mutations shown as spikes (the information is also mapped to the primary structure in **Figure 1A)** and selected antibody binding is shown schematically in **Figure 5E-G**. As expected there is correlation between the two, although the antibody centroids are more broadly spread across the RBD surface, in particular there are no mutations in the left flank epitope, where a significant number of antibodies bind (**Figure 5A**). These antibodies can neutralize in some assays, and confer protection (Barnes et al., 2020; Dejnirattisai et al., 2021a) and this cryptic epitope might therefore be an important target for therapeutic antibody applications and cross-protective vaccine antigen (Pinto et al., 2020). We demonstrate the continued binding of this class of antibodies below.

**Figure 5.**
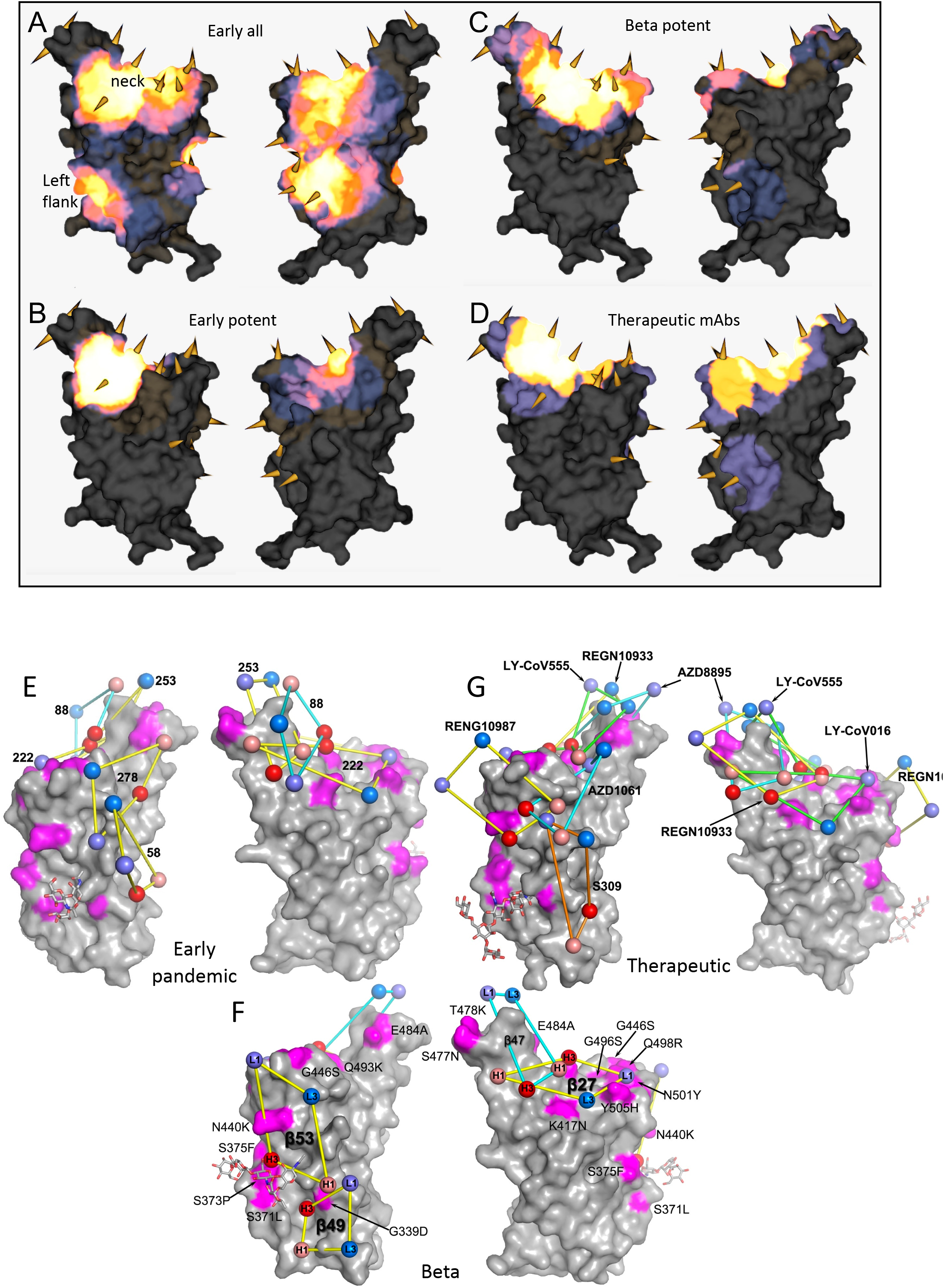
**Relative Antibody Contact**. RBD surface rendered in PyMOL exported and rendered in mabscape using iron heat colours (grey > straw > blue > glowing red > yellow > white) to indicate relative levels of antibody contact. Antibody contact is calculated for each surface vertex as the number of antibodies within a 10 Å radius by their known or predicted positions from earlier mapping studies (Dejnirattisai et al., 2021a; Liu et al., 2021b). Outward facing cones are placed at the nearest vertex to each mutated residue Calpha atom on the RBD surface. Drawn back and front views for (A) all RBD-reactive antibodies isolated from early pandemic or strongly neutralizing (< 100 ng/ml) (B) strongly neutralising antibodies isolated from Beta-infected sera (C) potent antibodies isolated from Beta infected cases (D) therapeutic antibodies for clinical use (from PDB: 7BEP, 6XDG, 7L7E, 7KMG, 7KMH). (A,B,C) Front (left) and back (right) views of the RBD drawn as a grey surface with Omicron changes highlighted in magenta and glycans drawn as sticks. (E) Outline footprints of a selection of early pandemic mAbs: 58, 88, 222, 253, 278 are shown by balls representing the centroid of interacting loops for LC (blues), HC (reds) and joined by yellow lines. (F) As for A, showing a selection of Beta antibodies:27, 47, 49, 53. Substituted residues in magenta are labelled. (G) As for A showing the footprints of a selection of commercial antibodies: REGN10933, REGN10987, S309, AZD1061, AZD8895, LY-CoV555, LY-CoV016.

Nineteen of the 20 most potent (FRNT50<100ng/ml) neutralising monoclonal antibodies we isolated from early pandemic cases mapped to the ACE2 binding site across the neck and shoulder epitopes of the RBD and 5 of these classified as public IGVH3-53 antibodies (Dejnirattisai et al., 2021a; Yuan et al., 2020). Mapping these onto the RBD surface (**Figure 5B**) shows that the centroids are highly concentrated in the neck region. IGVH3-53 mAbs were especially common in early pandemic responses and although their centroid is at the neck they are orientated such that their light chain CDRs interact with the right shoulder (**Figure S3**). Most IGVH3-53 mAbs are sensitive to the N501Y mutation, although some such as mAb 222 or Beta-27 can still neutralize 501Y containing viruses (Dejnirattisai et al., 2021b; Liu et al., 2021b). Omicron mutation Y505H makes a direct interaction with the L1 and L3 CDRs of mAb 222 (Dejnirattisai et al., 2021b). This, together with Q493R, which may also impact interaction with the H3 loop are likely responsible for the 12.6-fold reduction in the neutralization titre of mAb 222 (**Figure 4A, Table S1**).

MAb 253 as well as 55 and 165 are IGVH1-58 mAbs which bind an epitope shifted towards the left shoulder with H3 CDR contacting S477N and probably more importantly, Q493R may be disruptive, in this case approaching the H2 CDR, leading to the 19.3-fold reduction in neutralization of Omicron (**Figure S3**).

The neutralizing activity of mAbs 88, 316, and 384 is knocked out for Omicron (**Figure 4A, Table S1**), all interact with E484 (mAb 316 via H1 and H2) within the left shoulder epitope and the E484A mutation would be unfavourable. In the case of mAb 316, Q493 makes direct interactions with H1 and H3, thus Q493R will also likely be deleterious. Broadly neutralizing mAb 58 binds at the front of the RBD reaching towards the right flank in an area which is relatively clear of mutations and thus is unaffected (**Figure S3**). MAb 278 binds in a similar area but higher, the epitope comprising more of the right shoulder with L3 in contact with G446, and the G446S mutation in Omicron appears to knock out activity (**Figure S3**).

MAb 170 will be affected by Q493R and Q498R, which make direct interactions with L1 and H3 respectively (**Figure S3**). Q498R is between G496S and G446S (**Figure 2B**) and these substitutions may act in concert but since G446 is in proximity to H1 together these mutations knock out the activity of mAb 170 (**Figure 4A, Table S1**). The binding sites of selected potent antibodies including two which are cross-reactive against previous variants are shown in **Figure 5E**. The neutralization of all of these, with the exception of mAb58, are affected by the mutations in Omicron. To understand the resilience of nAb58 we determined the structure of a ternary complex of early pandemic RBD with Fabs for mAbs 58 and 158 (**Table S2**) confirming that its epitope includes no residues mutated in Omicron (**Figure S3**).

### Effect of Omicron mutations on Beta antibody responses

We have derived a panel of 27 potent Beta antibodies (FRNT <100 ng/ml) (Liu et al., 2021b) and this revealed a surprisingly skewed response with 18/27 potent antibodies targeting the Beta mutations: E484K, K417N and N501Y. This is seen in **Figure 5C**, where the focus on residues in the shoulders has spread the centroid patch out towards several Omicron mutation sites, this information is mapped to the primary structure in **Figure 1A** and a schematic of the binding of the four potent cross-reactive antibodies in **Figure 5F**. Whilst K417N and N501Y are conserved in Omicron, E484 is mutated to an alanine, which seems a likely escape mutation from either 484E (early pandemic/Alpha) or 484K (Beta).

Neutralization assays were performed against Omicron and show a complete loss of activity for 17/27 Beta mAb (**Figure 4B, Table S1**). Substantial reductions in neutralization titres were observed for many of the rest of the Beta panel, with Beta-22, 29, 40, 47, 53, 54, 55 and 56 able to neutralize Omicron with titres < 400ng/ml.

A large number of Beta mAbs target the 501Y mutation including a public antibody response mediated through IGVH4-39 (n=6) and the related IGVH4-30 (n=1) (Liu et al., 2021b) and many are likely to be sensitive to the numerous mutations in this region: N440K, G446S, Q493R, G496S, Q498R and Y505H. In total 11 antibodies make contact with 501Y; *Beta-**6, 10, 23, 24, 30,** 40, 54, 55, 56,* whilst Beta-22 and 29 bind epitopes dependent on 417N/T together with 501Y, (antibodies in italic are VH4-39 or 4-30 and the neutralization of Omicron for those in bold is completely knocked out). Beta MAbs targeting the back of the neck epitope (Beta- 22,29,30) will be affected, for example in the case of Beta-29, H1 makes extensive interactions with residues Q493, G496 and Y505 (**Figure S4**). Beta-44 binding to the left shoulder epitope has already been shown to be sensitive to T478K whilst the combination of S477N and T478K in Omicron is likely to be more deleterious. Interestingly several of the antibodies (Beta *40, 54, 55, 56* and 22, 29 (501Y 417N/T)) retain some activity and this is explained below with reference also to the structure of the Omicron RBD/Fab 55 complex, where we suggest that this may reflect Omicron escape from early pandemic rather than Beta responses.

Four Beta-mAbs have been shown to potently cross-neutralize all Alpha, Beta, Gamma and Delta variants (Liu et al., 2021b), their binding sites are shown in **Figure 5F**. Of these, Beta-27 is a VH3-53 antibody, which contacts Q493 and Y505 in a similar way to mAb222 and shows reduced neutralization of Omicron (**Figure 4B, Table S1**). Beta-47, a VH1-58 antibody, binding towards the left shoulder epitope is not sensitive to T478K mutation, but having contacts with S477 and Q493 likely lead to the observed reduction in neutralization of Omicron.

Beta-49 and 50 similarly bind to the right flank epitope and activity is knocked out on Omicron (**Figure 4B, Table S1**). Beta-49 and 50 both belong to the IGVH1-69 gene family, H3 lies directly above RDB G399 and would be expected to clash with G399D. Beta-53 also binds to the right flank with H1 contacting residue 339 and likely clashing with G339D and L1 will likely contact G446S leading to the observed two log reduction in Omicron neutralization compared to Beta (**Figure S4**).

### Effect of Omicron mutations on current antibody therapeutics

A number of individual antibodies or, cocktails of potently neutralizing antibodies (usually combinations recognizing different epitopes to avoid the likelihood of escape (Sun et al., 2021)) have been licensed for use and the aggregate of their binding shown schematically in **Figure 5G**, which illustrates the strong correlation of aggregate binding with points of mutation and this is shown mapped to the primary structure in **Figure 1A**. Neutralization of Omicron is markedly reduced in the majority of these antibodies (**Figure 4C, Table S1**). More specifically:

*Regeneron 10987 and 10933*: Regeneron 10933 (Weinreich et al., 2021) binds the back of the left shoulder and 10987 the right shoulder (**Figure 5G, S5**) activity of both is knocked out on Omicron (**Figure 4C**). It has already been shown that REGN 10933 was unable to effectively neutralize Beta being sensitive to E484K (Zhou et al., 2021). REGN10933 H2 also contacts Q493 and neutralizing activity to Omicron is almost completely lost. REGN10987 H3 has direct contact with N440 and H2 with G446 which cause complete neutralization loss (**Figure S5**).

*Vir S309*: S309 (Dejnirattisai et al., 2021a; Pinto et al., 2020; Sun and Ho, 2020) binds on the right flank with H3 contacting G339 and N343 glycans the latter close to the Serine 371, 373 and 375 mutations (**Figure 5G, S5**). S309 neutralization of Omicron is only reduced 5.5-fold compared to Victoria. The modest effect of these mutations was confirmed by surface plasmon resonance (SPR) measurements of S309 binding to both RBDs (**Figure S6A**).

*AZD8895 & AZD1061*: AZD8895, which binds the back of the left shoulder, activity on Omicron is reduced 197-fold compared to Victoria, and AZD1061, which is reduced 428-fold compared to Victoria, binds the front of the right shoulder (**Figures 5G, S5**). AZD1061 is affected due to contacts with the G446 loop (L2 and H3). AZD8895 a VH1-58 antibody like 253 & Beta-47, contacts S477 (H3) & Q493 (H2) and like 253 and Beta-47 is compromised. The activity of AZD7442 (a combination of AZD8895 and AZD1061) still maintains neutralizing activity against Omicron, although reduced 37.1-fold compared to Victoria.

*LY-CoV016 and 555*: Activity of both antibodies on Omicron is knocked out. LY-CoV016 is a VH3-53 antibody and has extensive interactions with N501 and Y505 via L1 and L3 making it vulnerable to mutations at these residues (**Figure 5G, S5**). LY-CoV555 (Sun and Ho, 2020) is sensitive to the E484K mutation in delta (Liu et al., 2021a) and also contacts Q493.

#### ADG 10, 20, 30

All of the Adagio antibodies suffer considerable loss of activity against Omicron (**Figure 4C**). Activity of ADG10 and ADG30 are completely lost whilst the activity of ADG20 on Omicron is reduced 276-fold

### Effect on RBD/ACE2 Interaction

Fitness of a virus can stem from higher infectivity or evasion of the immune system. One way to identify mutations that increase binding affinity is by selection using a randomly mutated RBD displayed on the yeast surface for ACE2 binding. Mutations fixed for higher affinity binding included N501Y, E484K, S477N and most prominently, Q498R (**Figure 6A**) (Zahradnik et al., 2021b). Interestingly, Q498R was selected only at later stages. This is explained by the 2-fold reduction in affinity as a single mutation (**Figure 6A**). However, in combination with the N501Y mutation, the affinity is increased 26-fold, more than any other mutation analyzed. Adding to this the S477N mutation, one obtains a 37-fold increase in binding (**Figure 6B**). These three mutations, selected through *in vitro* evolution, were found together for the first time in the Omicron variant.

**Figure 6.**
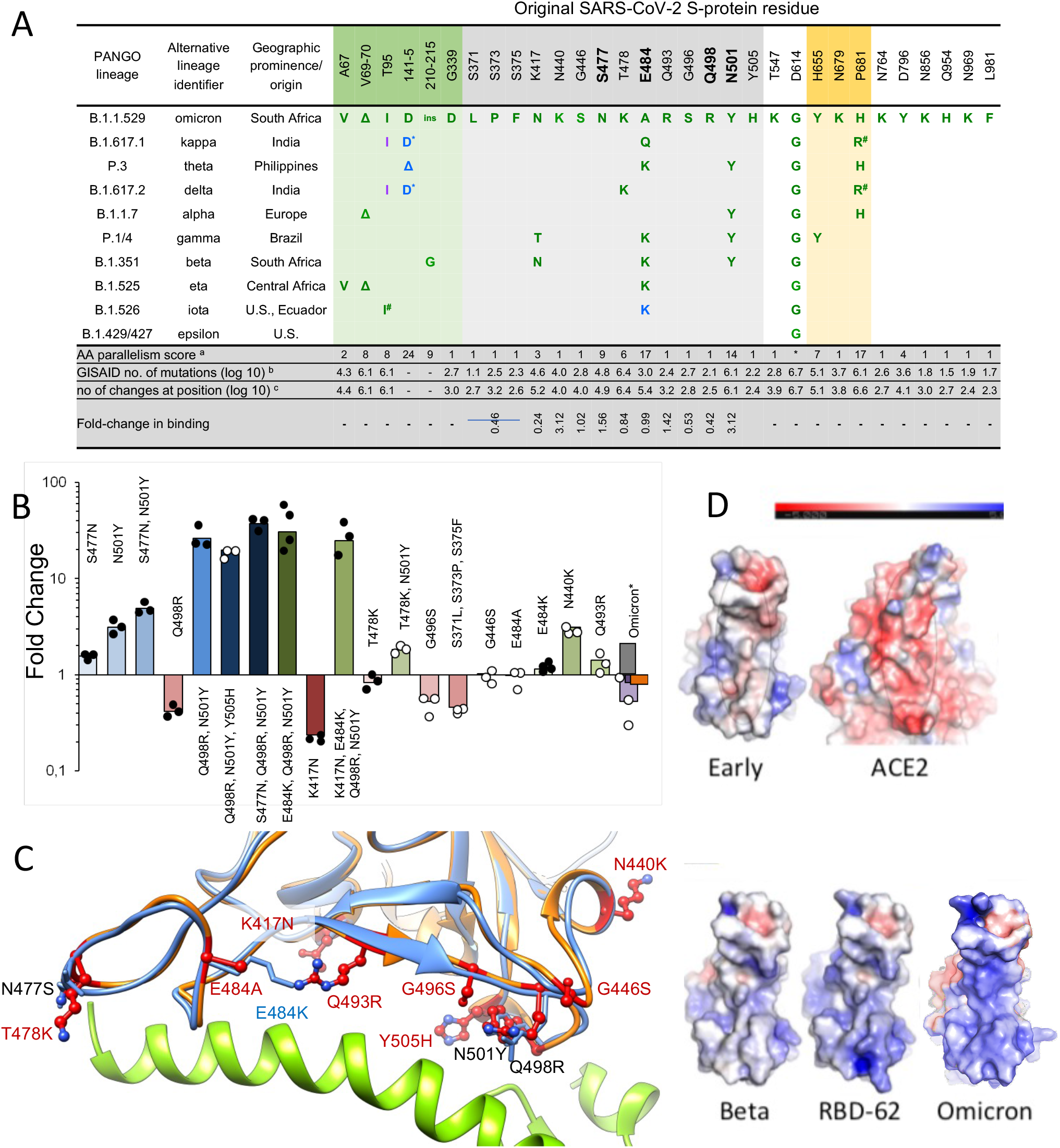
Affinity driving mutations in Omicron RBD have previously been identified by in vitro evolution for tighter binding. (A) Analysis of the occurrence and prevalence of Omicronvariant mutations. The background is coloured according to S-protein functional domains: NTD domain (AA 12 – 306; orange), RBD domain (AA 318 – 514; green), Furin cleavage site and its proximity (AA 655 – 701; blue). The four positions critical for the high affinity of RBD- 62 are highlighted in bold. Mutation frequencies within individual lineages are denoted in green (100-75 %), blue (75-50 %) and magenta (50-25 %). Information about the distribution and frequency of S-protein mutations and the spatiotemporal characterization of SARS-CoV- 2 lineages was retrieved from www.outbreak.info (Mullen et al., 2020) and Gisaid database (Elbe and Buckland-Merrett, 2017).* Same evolutionary origin, a Number of evolutionary non-related lineages with given or similar mutation (Zahradnik et al., 2021c), b log(10) number of the observed Omicron mutation at the given position as determined on 14.11.2021, c same as b but total log(10) number of changes at the given position. d fold- change in binding as determined by yeast-surface display. (B) Comparison of fold change in binding affinity among selected mutations and their combinations as determined by titrating ACE2 on yeast surface displayed RBD mutations. Values are fold-change relative to the original strain. For Omicron, yeast titration is denoted in violet, SPR (this study) is black, SPR as determined in (Cameroni et al., 2021) is grey and ELISA as determined in (Schubert et al., 2021) is in orange. Data denoted by black dots have been reported previously (Zahradnik et al., 2021b). (C) RBD-62 (blue)/ACE2 (green) structure (PDB: 7BH9) overlaid on Omicron RBD structure (orange) as determined bound to Beta 55. All Omicron mutations are shown, overlaid on relevant RBD-62 mutations. (D) Electrostatic potential surface depictions calculated using the APBS plugin in PyMol for left to right: early pandemic Victoria RBD showing the ACE2 interacting surface, ACE2 showing surface that binds the RBD, Beta RBD ACE2 interacting surface, RBD-62 ACE2 interacting surface, Omicron RBD ACE2 interacting surface. Blue is positive and red negative potential (scale bar shown above).

We measured the affinity of Omicron RBD for ACE2 and perhaps surprisingly, the affinity was on a par with that of the early virus, 8 nM and 7 nM respectively (these and binding affinities for a variety of other RBDs are shown in **Figure S6A**), implying that the increased affinity imparted by S477N, Q498R and N501Y are being offset by other mutations in the ACE2 footprint. We measured the affinities of the other single mutations in the ACE2 binding footprint of Omicron, shown in **Figure 6B, C**, and they provide a rationale for this. T478K in the presence of N501Y decreased the positive effect of the later by 2-fold. Y505H reduces binding of Q498R, N501Y double mutant by 50%. G496S and the triple-mutation S371L, S373P and S375F reduce binding by 2-fold and 2.2-fold respectively. E484A (instead of the Lys found other variants, **Figure 6A**) was neutral. While K417N (found in the Beta variant) on its own decreases binding substantially, the effect on binding when combined with other mutations is smaller **(Figure 6B)**. Two single-mutations found specifically in Omicron, Q493R and N440K did increase binding, probably due to increasing the electrostatic complementarity between ACE2 (negatively charged) and the RBD (positively charged) **(Figure 6D)**.

Comparing the RBD-62/ACE2 structure (PDB:7BH9) to that of Omicron RBD bound to Beta 55 antibody (described below, see **Table S2, Figure 6C)**, shows high similarity, with an RMSD of 0.55 Å over 139 residues. Importantly, the locations of the binding enhancing mutations 477N, 498R and 501Y are conserved between the two, despite the RBD-62 bound to ACE2, while Omicron RBD is not. This shows that these residues are pre-arranged for tight binding, implying low entropic penalty of binding.

### Antigenic Cartography

We have used the matrix of neutralization data generated in **Figure 3** to place Omicron on an antigenic map. We used a method similar to that developed for analysis of the Delta variant (Liu et al., 2021a), where we model individual viruses independently and allow for serum specific scaling of the responses (Methods). This simple model works well, the measured and modelled responses are shown in **Figure 7A,B** (with 1600 observations and 215 parameters the residual error is 9.1 %). The results are well described in three dimensions, see Video S1, and are shown projected into two dimensions in **Figure 7C,D**. It will be seen that the previous variants are placed in a planetary band around a central point, with Delta opposed to Beta and Gamma, Omicron is displaced a large distance out of this plane, almost on a line drawn from the central point perpendicular to the planetary band, illustrating vividly how Omicron dramatically expands our view of the antigenic landscape of SARS-CoV-2.

**Figure 7.**
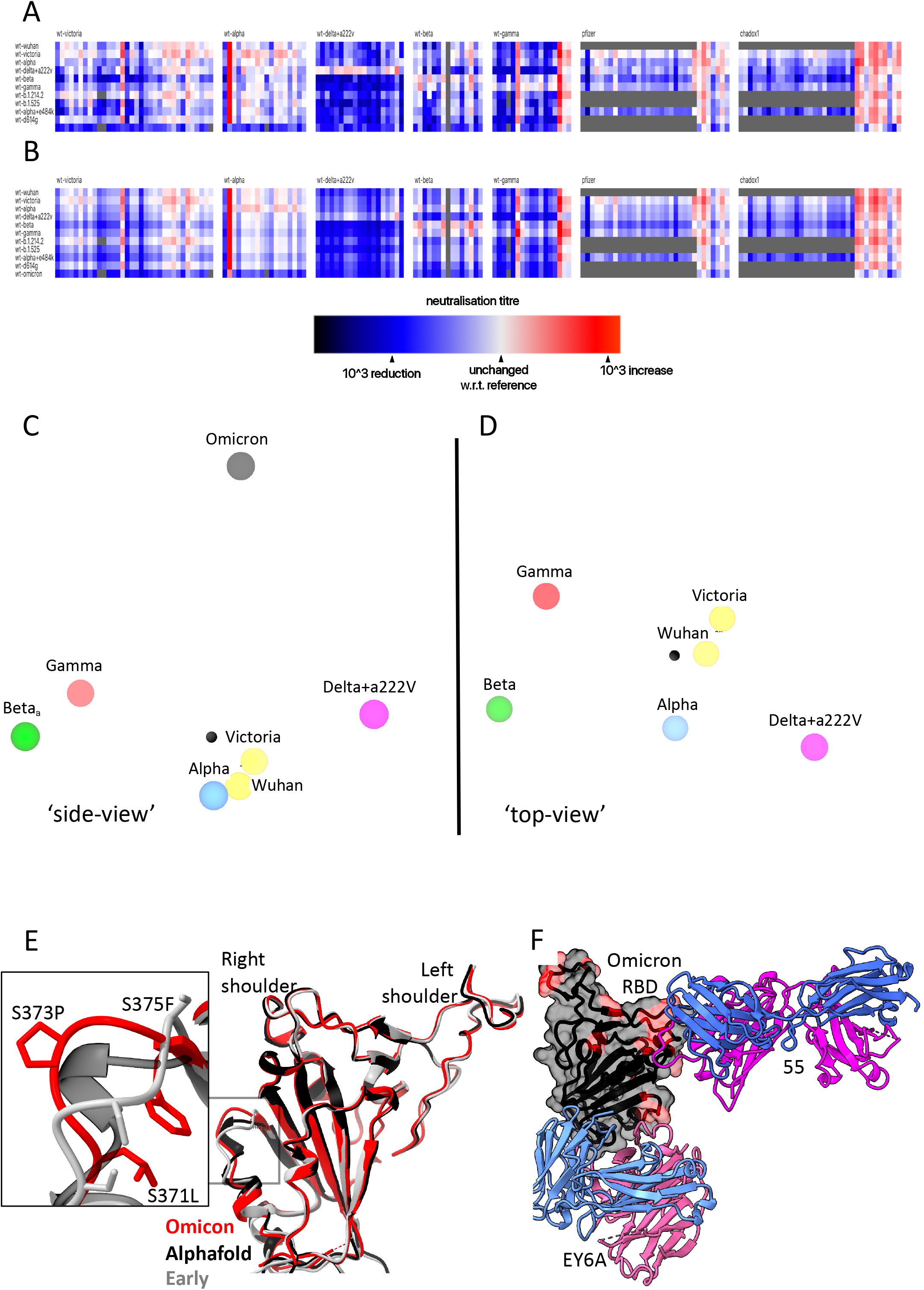
Antigenic map from neutralization data for omicron. (A) neutralization data (log titres) showing sera as columns against challenge variants as rows. Sera are grouped into blocks according to the eliciting variant. The reference neutralization titre for each block is calculated as the average of all self-challenge titres, i.e. when challenged with the same variant as which elicited the serum. In the case of vaccine sera this was taken as the average of all best neutralisation titres. Colours within a single block therefore express the relative neutralization titre with respect to this reference. (B) shows an example of the equivalent model generated from one run of antigenic map refinement using the same reference offsets as calculated for (A). (C) shows three-dimensional view of the antigenic map for variants of concern. The distance between two points corresponds to the drop-off in neutralisation titre used to generate value for (B). (D) same antigenic space as (C), but rotated to look down from the point of view of the Omicron position on the remaining variants. (E) Overlay of the X-ray structure of Omicron (red) on the early pandemic (Wuhan) RBD (grey) and the predicted model of the Omicron RBD in black, drawn as cartoons. The structural change effected by the S371L, S373P and S375F mutations is shown enlarged see inset. (F) X-ray structure of ternary complex of Omicron RBD with Beta 55 and EY6A Fabs. The Omicron RBD is shown as a grey semi-transparent surface with the mutated residues in magenta. The Fabs are drawn as cartoons, heavy chain in magenta and light chain in blue.

### The structural impact of the numerous mutations in S

To quickly gain insight into the possible structural impact of the numerous RBD mutations we used Alphafold2 (Jumper et al., 2021) to predict the Omicron RBD structure (**Figure S1B**). The top two ranked solutions, were essentially identical to each other and to the early pandemic reference structure (for residues 334 to 528 the RMSD for the 195 Ca atoms was 0.71 Å between the top ranked Alphafold2 prediction and the early pandemic structure). There was one region of significant, but still relatively minor, difference in the region of the triple serine mutation 371-375 (numbering for the early pandemic virus), on the right flank (**Figure S1B**). These mutations S371L, S373P, S375F all change from small flexible polar serine residues to bulkier, less flexible hydrophobic residues. Interestingly, all the Omicron S mutations involve single codon changes apart from S371L which requires two changes from TCC to CTC indicative of an underlying strong selection pressure and biological function to this change. The structure is not markedly altered however, it is exactly this region of the structure that undergoes a large conformation change when lipid is bound into the pocket. The serine rich loop opens, allowing the attached helix to swing out and open the pocket for lipid binding.

We suggest that increased rigidity and the entropic penalty of exposing hydrophobic residues may disfavour lipid binding to Omicron, which is likely to have an effect on the properties of the virus, explaining the selection of this apparently concerted series of changes.

To test these predictions, we determined the high-resolution crystal structure of the Omicron RBD domain in complex with two Fabs: Beta55 and EY6A, **Figure S6B, Table S2** (Liu et al., 2021b; Zhou et al., 2020). The RBD structure is very close to that observed in early pandemic viruses (RMSD 0.9 Å for 187 C*α*) and, as predicted by Alphafold2 the only significant change in the vicinity of the set of 3 serine mutations at residues 371-375 (**Figure 7E**). The rearrangement in this region is essentially an amplified version of that predicted, suggesting that algorithms such as Alphafold2 have some value in predicting the effect of dense patterns of mutations such as those seen in the Omicron RBD.

The binding of EY6A to the left flank of the RBD is essentially unchanged from that observed previously (Zhou et al., 2020) (KD 7.8 and 6.8 nM for early pandemic and Omicron RBDs respectively) (**Figure S6A, 7F**), in line with this cryptic epitope, which is highly conserved for functional reasons, being a good target for broadly neutralising therapeutic antibodies.

Beta55, as predicted earlier(Liu et al., 2021b), binds to the right shoulder, around residue 501. Interestingly, the epitope includes several residues mutated in Omicron from the early pandemic virus (including Q498R, N501Y, Y505H) (**Figure 7E, S6B, S6C**). It is remarkable that despite these significant changes, neutralization is relatively little affected. The neutralisation result was confirmed by measurements of the binding affinity, 177 nM and 204 pM for the early pandemic and Omicron RBDs respectively (**Figure S6A**). To confirm the structural basis. we also determined the crystal structure of an analogous ternary complex formed with early pandemic RBD (Table S2), as expected the details of the interaction are essentially identical. If we extend the analysis of the 501Y targeting antibodies by comparing the structures of Beta-6, 24, 40 and 54, we find subtle explanations, thus Beta-24 and some others, are knocked out due to a clash with CDR-L1 created by the Q493R mutation (**Figure S6D**), whereas for antibodies Beta-40, 54 and 55 this mutation can be accommodated. In addition, the Q498R mutation may create a hydrogen bond in Beta-40 or a salt bridge in Beta-54 to CDR-H3, which may compensate for the loss of binding affinity due to changes around residue 501 (**Figure S6E**). Thus, the surprising resilience of several of the 501Y targeting antibodies may be because the mutated residues in this region are not ‘hot-spots’ of interaction and mutations can sometimes be accommodated without significant impact on affinity. This may suggest that a major driver for evolution was the less 501-focussed responses to early viruses.

## Discussion

The first 4 Omicron sequences were deposited on 24^th^ November 2021. Within days distant international spread was seen and it is causing great concern due to its high transmissibility and ability to infect previously exposed or vaccinated individuals. Only three weeks after the virus was first detected in the UK Omicron cases outnumber Delta in London and the number of daily new cases in the UK is larger than at any previous time in the pandemic. Over the next weeks disease severity will become clearer.

The density of mutational changes (including deletions and insertions) found in Omicron S is extraordinary, being more than three times that observed in previous variants. Within S, as observed for other variants, the NTD, RBD and the furin cleavage site region are hotspots for mutation (Zahradnik et al., 2021b) and within the RBD, mutations are concentrated on the ACE2 interacting surface and the right flank. There seem to be two main drivers for the evolution of the RBD; increasing affinity to ACE2 and escape from the antibody response, which are coupled in Omicron.

Most potent neutralizing antibodies bind on, or in close proximity to the ACE2 footprint (neck and shoulder epitopes) and block interaction of S with ACE2, thereby preventing viral attachment to the host cell. There are two other classes of potent neutralizing mAbs, firstly antibodies binding in close proximity to the N343 glycan (right flank epitope) exemplified by Vir S309(Pinto et al., 2020) which include the Beta 49, 50 and 53 antibodies (Liu et al., 2021b) used in our analysis. These mAbs bind distant from the ACE2 binding site, do not block ACE2 interaction and their mechanism of action may be to destabilize the S trimer. Finally, antibodies binding to the supersite on the NTD can also be potently neutralizing although the mechanism of action of NTD antibodies remains obscure (Cerutti et al., 2021; Chi et al., 2020; Dejnirattisai et al., 2021a). Multiple mutations at all three of these sites: the receptor binding site, proximal to N343 glycan and NTD are found in Omicron, and lead to substantial reduction in neutralization titres for naturally immune or vaccine sera, with many showing complete failure of neutralization. This together with the widespread failure of potent mAb to neutralize Omicron point to a driver of immune evasion for their evolution.

The left flank epitope, which is not mutated in Omicron, in contrast is used by antibodies that do not block ACE2 binding, are not classed as potently neutralising in many assays and yet have been shown to be protective in animal models (Barnes et al., 2020; Dejnirattisai et al., 2021a; Hastie et al., 2021; Zhou et al., 2020). This strongly conserved epitope is inaccessible in most conformations of the RBD and binding is proposed to destabilise the S trimer (Benton et al., 2020; Huo et al., 2020; Zhou et al., 2020). It is possible that structural constraints on this epitope may render escape more problematic and this may therefore be an important target. Here we demonstrate structurally and by affinity measurements that the epitope is conserved and antibody binding essentially unchanged.

Following repeated rounds of selection by yeast display for high ACE2 affinity, RBD-62 (I358F, V445K, N460K, I468T, T470M, S477N, E484K, Q498R, N501Y) emerged as the highest affinity clone with a 1000-fold increase in affinity for ACE2 from 17 nM for Wuhan RBD to 16 pM for RBD-62. It is striking that the key contributors for the high affinity of RBD-62 are present in Omicron. Interestingly, the combination of mutations K417M, E484K, Q493R, Q498R and N501Y also emerged after 30 passages in mouse lungs (Wong et al., 2021). This mouse-adapted virus was highly virulent and caused a more severe disease. The appearance of E484K, Q493H/R, Q498R and N501Y in yeast display and mouse adaptation experiments is a strong indication that the tighter binding to ACE2 also facilitates more efficient transmission.

However, in Omicron. the virus opts for a different strategy, affinity to ACE2 is not increased for Omicron-RBD. Since mutations S477N, Q498R and N501Y are likely to increase ACE2 affinity by 37-fold, we hypothesise that these changes, also found in RBD-62, serve to anchor the RBD to ACE2, leaving the rest of the receptor binding motif more freedom to develop further mutations, including those that reduce ACE2 affinity, in the quest to evade the neutralizing antibody response. Indeed, K417N, T478K, G496S, Y505H and the triple S371L, S373P, S375F reduce affinity to ACE2, while driving immune evasion. All this is achieved with very minimal structural changes in the isolated Omicron RBD (**Figure 7E** ).

These observations provide a valuable lesson on the plasticity of protein-protein binding sites, maintaining nM binding affinity (Cohen-Khait and Schreiber, 2016). Thus, whilst the extreme concentration of potent neutralizing antibodies around the 25 amino acid receptor footprint of ACE2 suggest that this would be an Achilles heel for SARS-CoV-2, with ACE2 placing constraints on its variability (with receptor binding sites therefore sometimes hidden (Rossmann et al., 1985)). In practice, the extraordinary plasticity of this site to absorb mutational change, whilst retaining affinity for ACE2 is a potent weapon to evade the antibody response. Such camouflage of receptor binding sites has been observed before (see for example (Acharya et al., 1989), but it seems that by acquiring a lock on the ACE2 receptor at one point, through 498 and associated mutations many other less energetically favorable changes can be tolerated, fueling antigenic escape . Thus, by mutating the receptor binding site, the virus can modulate ACE2 affinity and potentially transmissibility, whist at the same time evading the antibody response.

How Omicron evolved is currently under debate. The results presented here show that immune evasion is a primary driver in its evolution, sacrificing the affinity enhancing mutations for optimizing immune evading mutations. This could for instance happen by a combination of a single immunocompromised individual which further evolved in rural, unmonitored populations (Clark et al., 2021). Virus evolution has been previously observed in chronically infected HIV+ individuals and other immunocompromised cases leading to the expression of the N501Y, E484K and K417T mutations (Cele et al., 2021; Karim et al., 2021; Kemp et al., 2021). What seems beyond doubt from the ratio of nonsynonymous to synonymous mutations (only one synonymous mutation in all of S) is that the evolution has been driven by strong selective pressure on S. It has been predicted that increasing immunity by natural infection or vaccination will increase the selective pressure to find a susceptible host, either by increased transmissibility or antibody evasion, it appears that Omicron has achieved both of these goals.

In addition to changes in the ACE2 footprint Omicron RBD possesses a triplet of changes from serines to more bulky, hydrophobic residues, a motif not found in any of the other Sarbecoviruses. This introduces some structural changes and may led to loss of ability to form the lipid binding pocket which might normally aid release of the virus from infected cells. Since one of these mutations requires a double change in the codon it is likely that this effect is significant and it is conceivable that it acts in synergy with the change at residue 498, perhaps explaining why, even in the context of N501Y, present in Alpha, Beta, Gamma and other minor variants, this mutation has not established itself earlier.

For most mAbs the changes in interaction are so severe that activity is completely lost or severely impaired. This also extends to the set of mAbs developed for clinical use, the activity of most is lost, AZD8895 and ADG20 activity is substantially reduced while the activity of Vir S309 is more modestly reduced.

Omicron has now got a foothold in many countries, in the UK it has an estimated doubling time of 2.5 days, 2 doses of vaccine appear to give low protection from infection, while 3 doses give better protection. There is considerable concern that Omicron will rapidly replace Delta and cause a large and sharp peak of infection in early 2022. It is likely that substantial increased transmissibility and immune evasion are contributing to the explosive rise in Omicron infections.

We have previously compared the neutralization of early pandemic SARS-CoV-2 strains, Alpha, Beta, Gamma and Delta using serum obtained early during the early pandemic or from Alpha, Beta or Gamma infected individuals. This allowed us to build a crude antigenic map of the SARS-CoV-2 sero-complex (Liu et al., 2021a). Early pandemic viruses and Alpha sit close to the centre, whilst Beta/Gamma diverge in one direction and Delta in the opposite direction, meaning that Beta/Gamma serum poorly neutralize Delta and vice versa. Unsurprisingly, when we recalculate the map including Omicron data we find that Omicron occupies the most antigenically distant position on the map, being almost orthogonal to the plane containing the Beta, Gamma and Delta.

At present, the only option to control the spread of Omicron, barring social distancing and mask wearing, is to pursue vaccination with Wuhan containing antigen, to boost the response to sufficiently high titres to provide some protection. However, the antigenic distance of Omicron may mandate the development of vaccines against this strain. There will then be a question of how to use these vaccines; vaccination with Omicron will likely give good protection against Omicron, but will not give good protection against other strains. It seems possible therefore that Omicron may cause a shift from the current monovalent vaccines containing Wuhan S to multivalent vaccines containing an antigen such as Wuhan or Alpha at the centre of the antigenic map and Omicron or other S genes at the extreme peripheries of the map, similar to the polyvalent strategies used in influenza vaccines.

In summary, we have presented data showing that the huge number of mutational changes present in Omicron lead to a substantial knock down of neutralizing capacity of immune serum and failure of mAb. This appears to lead to a fall in vaccine effectiveness, but it is unlikely that vaccines will completely fail and it is hoped that although vaccine breakthroughs will occur, protection from severe disease will be maintained, perhaps by T cells. It is likely that the vaccine induced T cell response to SARS-CoV-2 will be less affected than the antibody response. Third dose vaccine boosters substantially raise neutralization titres to Omicron and are, in countries such as the UK, the mainstay of the response to Omicron. Widespread vaccine breakthrough may mandate the production of a vaccine tailored to Omicron and failure of monoclonal antibodies may likewise lead to the generation of second generation mAbs targeting Omicron.

A question asked after the appearance of each new variant is whether SARS-CoV-2 has reached its limit for evolution. Analysing the mutations in Omicron shows that, except for S371L, all other mutations required only single-nucleotide changes. Also, most changes observed in the *in vitro* RBD-62 evolution study constituted single-nucleotide mutations. Two-nucleotide mutations and epistatic mutations are more difficult to reach, but open up vast untapped potentials for future variants. To avoid this, it is imperative to control the SARS-CoV-2 pandemic to reduce the number of infected people through vaccination and other measures globally.

## Limitations

The neutralization assays presented in this paper are performed *in vitro*, they do not therefore fully quantify the antibody response *in vivo* where complement and antibody dependent cell mediated cytotoxicity may contribute to virus control. Evasion of the antibody response may allow reinfection with Omicron, but the role of the T cell response, not measured here, is likely to contribute to the control of infection and moderate severe disease.

## Supporting information

Video of antigenic map of SARS-CoV-2 variants

## Acknowledgements

This work was supported by the Chinese Academy of Medical Sciences (CAMS) Innovation Fund for Medical Science (CIFMS), China (grant number: 2018-I2M-2-002) to D.I.S. and G.R.S. We are also grateful for support from Schmidt Futures, the Red Avenue Foundation and the Oak Foundation. H.M.E.D. and J.Ren are supported by the Wellcome Trust (101122/Z/13/Z), D.I.S. and E.E.F. by the UKRI MRC (MR/N00065X/1). G.S and J.Z were supported by the Israel Science Foundation (grant no. 3814/19) and O.A and S.K (grant no. 3729/20) within the KillCorona—Curbing Coronavirus Research Program and by the Ben B. and Joyce E. Eisenberg Foundation. D.I.S. and G.R.S. are Jenner Investigators. This is a contribution from the UK Instruct-ERIC Centre. AJM is an NIHR-supported Academic Clinical Lecturer. The convalescent sampling was supported by the Medical Research Council [grant MC_PC_19059] (awarded to the ISARIC-4C consortium) (with a full contributor list available at https://isaric4c.net/about/authors/) and the National Institutes for Health and Oxford Biomedical Research Centre and an Oxfordshire Health Services Research Committee grant to AJM. OPTIC Consortium: Christopher Conlon, Alexandra Deeks, John Frater, Lisa Frending, Siobhan Gardiner, Anni Jämsén, Katie Jeffery, Tom Malone, Eloise Phillips, Lucy Rothwell, Lizzie Stafford. The Wellcome Centre for Human Genetics is supported by the Wellcome Trust (grant 090532/Z/09/Z). The computational aspects of this research were supported by the Wellcome Trust Core Award Grant Number 203141/Z/16/Z and the NIHR Oxford BRC. The Oxford Vaccine work was supported by UK Research and Innovation, Coalition for Epidemic Preparedness Innovations, National Institute for Health Research (NIHR), NIHR Oxford Biomedical Research Centre, Thames Valley and South Midland’s NIHR Clinical Research Network. We thank the Oxford Protective T-cell Immunology for COVID-19 (OPTIC) Clinical team for participant sample collection and the Oxford Immunology Network Covid-19 Response T cell Consortium for laboratory support. We acknowledge the rapid sharing of Victoria, B.1.1.7 and B.1.351 which was isolated by scientists within the National Infection Service at PHE Porton Down, and the B.1.617.2 virus was kindly provided Wendy Barclay and Thushan De Silva. We thank The Secretariat of National Surveillance, Ministry of Health Brazil for assistance in obtaining P.1 samples. This work was supported by the UK Department of Health and Social Care as part of the PITCH (Protective Immunity from T cells to Covid-19 in Health workers) Consortium, the UK Coronavirus Immunology Consortium (UK-CIC) and the Huo Family Foundation. EB and PK are NIHR Senior Investigators and PK is funded by WT109965MA and NIH (U19 I082360). SJD is funded by an NIHR Global Research Professorship (NIHR300791). DS is an NIHR Academic Clinical Fellow. The team at the University of Witwatersrand were supported by The Bill & Melinda Gates Foundation (grant number INV- 016202). The views expressed in this article are those of the authors and not necessarily those of the National Health Service (NHS), the Department of Health and Social Care (DHSC), the National Institutes for Health Research (NIHR), the Medical Research Council (MRC) or Public Health, England.

## Author Information

These authors contributed equally: W.D., J.H., D.Z., J.Z., P.S., C.L.

## Contributions

J.H., J.Z., G.S., S.K. and O.A. performed interaction affinity analyses. D.Z., J.R., N.G.P., M.A.W. and D.R.H. prepared the crystals, enabled and performed X-ray data collection. J.R., E.E.F., H.M.E.D. M.B. and D.I.S. analysed the structural results. G.R.S., J.H., J.M., P.S., D.Z., B.W., R.N., A.T., A.D. and C.L. prepared the RBDs, ACE2 and antibodies and, W.D. and P.S. performed neutralization assays. D.C., H.W., B.C., A.R.T., K-Y.A.H., T.K.T provided materials. H.M.G. wrote MABSCAPE and performed mapping and cluster analysis, including sequence and antigenic space analyses. S.A.C.C., F. G. N., V.N., F. N., C. F.D.C., P.C.R., A.P-C., M.M.S., A.J.M., E.B., S.J.D., S.A., D.S., A.A., S.A.J, C.D., S.A., D.P., A.B., N.O-D, D.K., D.J. P.K.A., M.V., P.J.M.O., J.K.B., M.G.S., A.J.P., P.K., M.W.C., T.L., A.F., T.R.,C.M.,T.M., N.S.,Z.D.,K.D.S, M.C.N., S.M. assisted with patient samples and vaccine trials. E.B., M.W.C., S.J.D., P.K. and D.S. conceived the study of vaccinated healthcare workers and oversaw the OPTIC Healthcare Worker study and sample collection/processing. V.B. performed molecular testing and sequencing, G.R.S. and D.I.S. conceived the study and wrote the initial manuscript draft with other authors providing editorial comments. All authors read and approved the manuscript.

---

Figure S1. (A) Number of sequenced mutations per position. The line shows the number of mutations per residue, for high to low along the spike protein. In green are mutations D614G, which is fixed from early virus evolution and position 498, which became dominant only in omicron. Red are for mutations in Omicron that were identified before in multiple linages and blue are mutations with Omicron being the only lineage. (B) Location of the S371L, S373P and S375F mutations in the context of the conformation change occurring on binding lipid. Cartoons of the apo (blue) and lipid bound (pink) early pandemic RBD are shown. The lipid is shown in red. Related to Figures 2 and 7E.

Figure S2. FRNT50 values for 7 cases of Delta infection before and after vaccination. Related to Figure 4.

Figure S3. **Binding modes of early pandemic mAbs and their contacts to Omicron mutation sites.** Fabs are drawn as ribbons with the heavy chains in red and light chains in blue, and RBDs as grey ribbon or surface representation with Omicron mutation sites highlighted in magenta. Side chains are shown as sticks, and hydrogen bonds as dashed lines. (A) Fab 58 dosen’t make any close contacts with the Omicron mutation sites. (B)-(F) Binding modes and contacts with Omicron mutation sites of Fabs 170, 222, 253, 278 and 316 respectively. Related to Figure 5.

Figure S4. **Binding modes of Beta mAbs and their contacts to Omicron mutation sites.** The drawing and colouring schemes are same as in Figure S2. These are structures of Beta-RBD/Beta-Fab complexes. (A) Beta-24 and (B) Beta-54, examples of Beta mAbs targeting the N501Y mutation site. (C) Beta-38, a representative of Beta mAbs targeting the E484K mutation site. (D) Beta-29, a K417N/T dependent Beta mAb. (E) Beta-44 binds at the top of left shoulder and is sensitive to T478K mutation. (F)-(I) Beta-27, 47, 49 and 53 respectively. These four Beta mAbs neutralise all the previous variants of concern as well as the early pandemic Wuhan strain. Related to Figure 5.

Figure S5. **Binding modes of the therapeutic mAbs and their contacts to Omicron mutation sites.** The drawing and colouring schemes are same as in Figure S2. (A) REGN10987 and REGN10933. (B) AZD8895 and AZD1061. (C) Vir S309. (D) LY-CoV016 and (E) LY-CoV555. Related to Figure 5.

Figure S6. **SPR measurement and Crystal structure of the Omicron RBD complexed with Beta-55 and EY6A Fabs.** (A) SPR measuremments. (B) Ternary complex of the Omicron-RBD (grey)/Beta-55 (HC red, LC blue)/EY6A (HC salmon, LC cyan). (C) Electron density map showing the density for the mutated residues at 446, 498, 501 and 505, and their interactions with the CDR-H3 of Beta-55. (D) and (E) Comparison of the slightly different binding mode of Beta-55 to Beta-24 (cyan in (D)) and Beta-40 (cyan in (E)), the close-up boxes show details of the interactions with Beta-24 and Beta-40 explaining the knock-out of Beta-24 and the resilience of Beta-40. Related to Figure 7.

## STAR Methods

### RESOURCE AVAILABILITY

#### Lead Contact

Resources, reagents and further information requirement should be forwarded to and will be responded by the Lead Contact, David I Stuart.

#### Materials Availability

Reagents generated in this study are available from the Lead Contact with a completed Materials Transfer Agreement.

### Data and Code Availability

The coordinates and structure factors of the crystallographic complexes are available from the PDB with accession codes (see **Table S2**). Mabscape is available from https://github.com/helenginn/mabscape, https://snapcraft.io/mabscape. The data that support the findings of this study are available from the corresponding authors on request.

### EXPERIMENTAL MODEL AND SUBJECT DETAILS

#### Viral stocks

SARS-CoV-2/human/AUS/VIC01/2020 (Caly et al., 2020), Alpha and Beta were provided by Public Health England, Gamma cultured from a throat swab from Brazil, Delta was a gift from Wendy Barclay and Thushan de Silva, from the UK G2P genotype to phenotype consortium and Omicron was grown from a positive throat swab (IRAS Project ID: 269573, Ethics Ref: 19/NW/0730. Briefly, VeroE6/TMPRSS2 cells (NIBSC) were maintained in Dulbecco’s Modified Eagle Medium (DMEM) high glucose supplemented with 1% fetal bovine serum, 2mM Glutamax, 100 IU/ml penicillin-streptomycin and 2.5ug/ml amphotericin B, at 37 °C in the presence of 5% CO2 before inoculation with 200ul of swab fluid. Cells were further maintained at 37°C with daily observations for cytopathic effect (CPE). Virus containing supernatant were clarified at 80% CPE by centrifugation at 3,000 r.p.m. at 4 °C before being stored at -80 °C in single-use aliquots. Viral titres were determined by a focus-forming assay on Vero CCL-81 cells (ATCC). Sequencing of the Omicron isolate shows the expected consensus S gene changes (A67V, Δ69-70, T95I, G142D/Δ143-145, Δ211/L212I, ins214EPE, G339D, S371L, S373P, S375F, K417N, N440K, G446S, S477N, T478K, E484A, Q493R, G496S, Q498R, N501Y, Y505H, T547K, D614G, H655Y, N679K, P681H, N764K, D796Y, N856K, Q954H, N969K, L981F), an intact furin cleavage site and a single additional mutation A701V. Cells were infected with the SARS-CoV-2 virus using an MOI of 0.0001.

Virus containing supernatant were harvested at 80% CPE and spun at 3000 rpm at 4 °C before storage at -80 °C. Viral titres were determined by a focus-forming assay on Vero cells. Victoria passage 5, Alpha passage 2 and Beta passage 4 stocks Gamma passage 1, Delta passage 3 and Omicron passage 1 were sequenced to verify that they contained the expected spike protein sequence and no changes to the furin cleavage sites.

#### Bacterial Strains and Cell Culture

Vero (ATCC CCL-81) and VeroE6/TMPRSS2 cells were cultured at 37 °C in Dulbecco’s Modified Eagle medium (DMEM) high glucose (Sigma-Aldrich) supplemented with 10% fetal bovine serum (FBS), 2 mM GlutaMAX (Gibco, 35050061) and 100 U/ml of penicillin–streptomycin. Human mAbs were expressed in HEK293T cells cultured in UltraDOMA PF Protein-free Medium (Cat# 12-727F, LONZA) at 37 °C with 5% CO2. HEK293T (ATCC CRL-11268) cells were cultured in DMEM high glucose (Sigma-Aldrich) supplemented with 10% FBS, 1% 100X Mem Neaa (Gibco) and 1% 100X L-Glutamine (Gibco) at 37 °C with 5% CO2. To express RBD, RBD variants and ACE2, HEK293T cells were cultured in DMEM high glucose (Sigma) supplemented with 2% FBS, 1% 100X Mem Neaa and 1% 100X L-Glutamine at 37 °C for transfection. Omicron RBD and human mAbs were also expressed in HEK293T (ATCC CRL-11268) cells cultured in FreeStyle 293 Expression Medium (ThermoFisher, 12338018) at 37 °C with 5% CO2. *E.coli DH5α* bacteria were used for transformation and large-scale preparation of plasmids. A single colony was picked and cultured in LB broth at 37 °C at 200 rpm in a shaker overnight.

#### Plasma from early pandemic and Alpha cases

Participants from the first wave of SARS-CoV2 in the U.K. and those sequence confirmed with B.1.1.7 lineage in December 2020 and February 2021 were recruited through three studies: Sepsis Immunomics [Oxford REC C, reference:19/SC/0296]), ISARIC/WHO Clinical Characterisation Protocol for Severe Emerging Infections [Oxford REC C, reference 13/SC/0149] and the Gastro-intestinal illness in Oxford: COVID sub study [Sheffield REC, reference: 16/YH/0247]. Diagnosis was confirmed through reporting of symptoms consistent with COVID-19 and a test positive for SARS-CoV-2 using reverse transcriptase polymerase chain reaction (RT-PCR) from an upper respiratory tract (nose/throat) swab tested in accredited laboratories. A blood sample was taken following consent at least 14 days after symptom onset. Clinical information including severity of disease (mild, severe or critical infection according to recommendations from the World Health Organisation) and times between symptom onset and sampling and age of participant was captured for all individuals at the time of sampling. Following heat inactivation of plasma/serum samples they were aliquoted so that no more than 3 freeze thaw cycles were performed for data generation.

#### Sera from Beta, Gamma and Delta infected cases

Beta and Delta samples from UK infected cases were collected under the “Innate and adaptive immunity against SARS-CoV-2 in healthcare worker family and household members” protocol affiliated to the Gastro-intestinal illness in Oxford: COVID sub study discussed above and approved by the University of Oxford Central University Research Ethics Committee. All individuals had sequence confirmed Beta/Delta infection or PCR-confirmed symptomatic disease occurring whilst in isolation and in direct contact with Beta/Delta sequence-confirmed cases. Additional Beta infected serum (sequence confirmed) was obtained from South Africa. At the time of swab collection patients signed an informed consent to consent for the collection of data and serial blood samples. The study was approved by the Human Research Ethics Committee of the University of the Witwatersrand (reference number 200313) and conducted in accordance with Good Clinical Practice guidelines. Gamma samples were provided by the International Reference Laboratory for Coronavirus at FIOCRUZ (WHO) as part of the national surveillance for coronavirus and had the approval of the FIOCRUZ ethical committee (CEP 4.128.241) to continuously receive and analyse samples of COVID-19 suspected cases for virological surveillance. Clinical samples were shared with Oxford University, UK under the MTA IOC FIOCRUZ 21-02.

#### Sera from Pfizer vaccinees

Pfizer vaccine serum was obtained from volunteers who had received either one or two doses of the BNT162b2 vaccine. Vaccinees were Health Care Workers, based at Oxford University Hospitals NHS Foundation Trust, not known to have prior infection with SARS-CoV-2 and were enrolled in the OPTIC Study as part of the Oxford Translational Gastrointestinal Unit GI Biobank Study 16/YH/0247 [research ethics committee (REC) at Yorkshire & The Humber – Sheffield] which has been amended for this purpose on 8 June 2020. The study was conducted according to the principles of the Declaration of Helsinki (2008) and the International Conference on Harmonization (ICH) Good Clinical Practice (GCP) guidelines. Written informed consent was obtained for all participants enrolled in the study. Participants were studied after receiving two doses of, and were sampled approximately 28 days (range 25-38), 180 days (range 178-221) and 270 days (range 243-273) after receiving two doses of Pfizer/BioNtech BNT162b2 mRNA Vaccine, 30 micrograms, administered intramuscularly after dilution (0.3 mL each), 17-28 days apart, then approximately 28 days (range 25-56) after receiving a third “booster dose of BNT162B2 vaccine. The mean age of vaccinees was 37 years (range 22-66), 21 male and 35 female.

#### AstraZeneca-Oxford vaccine study procedures and sample processing

Full details of the randomized controlled trial of ChAdOx1 nCoV-19 (AZD1222), were previously published (PMID: 33220855/PMID: 32702298). These studies were registered at ISRCTN (15281137 and 89951424) and ClinicalTrials.gov (NCT04324606 and NCT04400838). Written informed consent was obtained from all participants, and the trial is being done in accordance with the principles of the Declaration of Helsinki and Good Clinical Practice. The studies were sponsored by the University of Oxford (Oxford, UK) and approval obtained from a national ethics committee (South Central Berkshire Research Ethics Committee, reference 20/SC/0145 and 20/SC/0179) and a regulatory agency in the United Kingdom (the Medicines and Healthcare Products Regulatory Agency). An independent DSMB reviewed all interim safety reports. A copy of the protocols was included in previous publications (Folegatti et al., 2020).

Data from vaccinated volunteers who received two vaccinations are included in this paper. Vaccine doses were either 5 × 10^10^ viral particles (standard dose; SD/SD cohort n=21) or half dose as their first dose (low dose) and a standard dose as their second dose (LD/SD cohort n=4). The interval between first and second dose was in the range of 8-14 weeks. Blood samples were collected and serum separated on the day of vaccination and on pre-specified days after vaccination e.g. 14 and 28 days after boost.

### Method Details

#### Focus Reduction Neutralization Assay (FRNT)

The neutralization potential of Ab was measured using a Focus Reduction Neutralization Test (FRNT), where the reduction in the number of the infected foci is compared to a negative control well without antibody. Briefly, serially diluted Ab or plasma was mixed with SARS-CoV- 2 strain Victoria or P.1 and incubated for 1 hr at 37 °C. The mixtures were then transferred to 96-well, cell culture-treated, flat-bottom microplates containing confluent Vero cell monolayers in duplicate and incubated for a further 2 hrs followed by the addition of 1.5% semi-solid carboxymethyl cellulose (CMC) overlay medium to each well to limit virus diffusion. A focus forming assay was then performed by staining Vero cells with human anti-NP mAb (mAb206) followed by peroxidase-conjugated goat anti-human IgG (A0170; Sigma). Finally, the foci (infected cells) approximately 100 per well in the absence of antibodies, were visualized by adding TrueBlue Peroxidase Substrate. Virus-infected cell foci were counted on the classic AID EliSpot reader using AID ELISpot software. The percentage of focus reduction was calculated and IC50 was determined using the probit program from the SPSS package.

#### DNA manipulations

Cloning was done by using a restriction-free approach(Peleg and Unger, 2014). Mutagenic megaprimers were PCR amplified (KAPA HiFi HotStart ReadyMix, Roche, Switzerland, cat. KK3605), purified by using NucleoSpin® Gel and PCR Clean-up kit (Nacherey-Nagel, Germany, REF 740609.50) and cloned into pJYDC1 (Adgene ID: 162458) (Zahradnik et al., 2021a). Parental pJYDC1 molecules were cleaved by DpnI treatment (1 h, NEB, USA, cat. R0176) and the reaction mixture was electroporated into E.coli Cloni® 10G cells (Lucigen, USA). The correctness of mutagenesis was verified by sequencing.

#### Yeast display binding assays

Plasmids (pJYDC1) with mutations were transformed (1 ug of DNA) by LiAc method (Gietz and Woods, 2006) into the EBY100 Saccharomyces cerevisiae and selected by growth on SD-W plates (Zahradnik et al., 2021a) for 48-72 h at 30°C. Grown single colonies were transferred to 1.0 ml liquid SD-CAA media, grown 24 h or 48 (RBD-Omicron) at 30°C (220 rpm), and 50 ul of the starter culture was used as inoculum (5 %) for the expression culture in 1/9 media (1 ml) supplemented with 1 nM DMSO solubilized bilirubin (Merck/Sigma-Aldrich cat. B4126). The expression continued in a shaking incubator for 24 h at 20°C (220 rpm). Aliquots of yeast expressed cells (100 ul, 3000 g, 3 min) were washed in ice-cold PBSB buffer (PBS with 1 g/L BSA) and resuspended in PBSB with a dilution series CF640-ACE2 (1 pM – 80 nM). The volume and incubation times were adjusted to limit the ligand depletion effect and enable equilibrium (Zahradnik et al., 2021b). After incubation, cells were washed in ice-cold PBSB buffer (PBS with 1 g/L BSA) passed through cell strainer nylon membrane (40 µM, SPL Life Sciences, Korea), and analyzed. The yeast expressing Omicron-RBD were expression labelled by primary anti-c-Myc 9E10 antibody (Biolegend, Cat. 626872) and secondary Anti-Mouse IgG(Fc specific)-FITC (Merck/Sigma-Aldrich, cat. F4143) antibodies. The signals for expression (FL1, eUnaG2 fluorophore, Ex. 498 nm, Em. 527 nm or FITC) and for binding (FL3, CF®640R dye-labeled ACE2) were recorded by S3e Cell Sorter (BioRad, USA). The standard non-cooperative Hill equation was fitted by nonlinear least-squares regression with two additional parameters using Python 3.7 (Starr et al., 2020; Zahradnik et al., 2021a).

#### Antigenic mapping

Antigenic mapping of omicron was carried out through an extension of a previous algorithm (Liu et al., 2021a). In short, coronavirus variants were assigned three-dimensional coordinates whereby the distance between two points indicates the base drop in neutralization titre. Each serum was assigned a strength parameter which provided a scalar offset to the logarithm of the neutralization titre. These parameters were refined to match predicted neutralization titres to observed values by taking an average of superimposed positions from 30 separate runs. The three-dimensional positions of the variants of concern: Victoria, Alpha, Beta, Gamma, Delta and Omicron were plotted for display.

#### Alphafold

Models of Omicron RBD and NTD were derived using AlphaFold 2.0.01 (Jumper et al., 2021) downloaded and installed on 11^th^ August 2021 in batch mode. For RBD predictions, 204 residues (P327-n529) were used as an input sequence while the NTD sequence input was from residues V1-S253. The max_release_date parameter was set to 28-11-2021 when the simulations were run such that template information was used for structure modelling. For all targets, five models were generated and all presets were kept the same.

#### Cloning of Spike and RBD

Expression plasmids of wild-type and Omicron spike and RBD were constructed encoding for human codon-optimized sequences from wild-type SARS-CoV-2 (MN908947) and Omicron (EPI_ISL_6640917). Wild-type Spike and RBD plasmids were constructed as described before (Dejnirattisai et al., 2021a). Spike and RBD fragments of Omicron were custom synthesized by GeneArt (Thermo Fisher Scientific GENEART) and cloned into pHLsec and pNEO vectors, respectively, as previously described (Dejnirattisai et al., 2021a; Supasa et al., 2021; Zhou et al., 2021). Both constructs were verified by Sanger sequencing after plasmid isolation using QIAGEN Miniprep kit (QIAGEN).

#### Protein production

Protein expression and purification were conducted as described previously (Dejnirattisai et al., 2021a; Zhou et al., 2020). Briefly, plasmids encoding proteins were transiently expressed in HEK293T (ATCC CRL-11268) cells. The conditioned medium was concentrated using a QuixStand benchtop system. His-tagged Omicron RBD were purified with a 5 mL HisTrap nickel column (GE Healthcare) and further polished using a Superdex 75 HiLoad 16/60 gel filtration column (GE Healthcare). Twin-strep tagged Omicron spike was purified with Strep-Tactin XT resin (IBA lifesciences). ∼4mg of ACE2 was mixed with homemade His-tagged 3C protease and DTT (final concentration 1mM). After incubated at 4 °C for one day, the sample was flown through a 5 mL HisTrap nickel column (GE Healthcare). His-tagged proteins were removed by the nickel column and purified ACE2 was harvested and concentrated.

#### IgG mAbs and Fab purification

To purify full length IgG mAbs, supernatants of mAb expression were collected and filtered by a vacuum filter system and loaded on protein A/G beads over night at 4 °C. Beads were washed with PBS three times and 0.1 M glycine pH 2.7 was used to elute IgG. The eluate was neutralized with Tris-HCl pH 8 buffer to make the final pH=7. The IgG concentration was determined by spectro-photometry and buffered exchanged into PBS.

To express and purify Fabs 158 and EY6A, heavy chain and light chain expression plasmids of Fab were co-transfected into HEK293T cells by PEI. After cells cultured for 5 days at 37°C with 5% CO2, culture supernatant was harvested and filtered using a 0.22 mm polyethersulfone filter. Fab 158 was purified using Strep-Tactin XT resin (IBA lifesciences) and Fab EY6A was purified with Ni-NTA column (GE HealthCare) and a Superdex 75 HiLoad 16/60 gel filtration column (GE Healthcare).

AstraZeneca and Regeneron antibodies were provided by AstraZeneca, Vir, Lilly and Adagio antibodies were provided by Adagio. For the antibodies heavy and light chains of the indicated antibodies were transiently transfected into 293Y cells and antibody purified from supernatant on protein A. Fab fragments of 58 and beta-55 were digested from purified IgGs with papain using a Pierce Fab Preparation Kit (Thermo Fisher), following the manufacturer’s protocol.

#### Surface Plasmon Resonance

The surface plasmon resonance experiments were performed using a Biacore T200 (GE Healthcare). All assays were performed with a running buffer of HBS-EP (Cytiva) at 25 °C. To determine the binding kinetics between the SARS-CoV-2 RBDs and ACE2 / monoclonal antibody (mAb), a Protein A sensor chip (Cytiva) was used. ACE2-Fc or mAb was immobilised onto the sample flow cell of the sensor chip. The reference flow cell was left blank. RBD was injected over the two flow cells at a range of five concentrations prepared by serial twofold dilutions, at a flow rate of 30 μl min^−1^ using a single-cycle kinetics programme. Running buffer was also injected using the same programme for background subtraction. All data were fitted to a 1:1 binding model using Biacore T200 Evaluation Software 3.1.

To determine the binding kinetics between the SARS-CoV-2 Spikes and ACE2, a CM5 sensor chip was used. The sensor chip was firstly activated by an injection of equal volume mix of EDC and NHS (Cytiva) at 20 uL/min for 300 s, followed by an injection of Spike sample at 20 ug/mL in 10 mM sodium acetate pH 5.0 (Cytiva) onto the sample flow cell of the sensor chip at 10 uL/min, and finally with an injection of 1.0 M Ethanolamine-HCl, pH 8.5 (Cytiva) at 20 uL/min for 180 s. The reference flow cell was left blank. ACE2 was injected over the two flow cells at a range of five concentrations prepared by serial twofold dilutions, at a flow rate of 30 μl min^−1^ using a single-cycle kinetics programme. Running buffer was also injected using the same programme for background subtraction. All data were fitted to a 1:1 binding model using Biacore T200 Evaluation Software 3.1.

#### Crystallization

Wuhan RBD was mixed with mAb-58 and mAb-158 Fabs, Wuhan or Omicron RBD was mixed with EY6A and beta-55 Fabs, in a 1:1:1 molar ratio, with a final concentration of 7, 7 and 3 mg ml^-1^. These complexes were separately incubated at room temperature for 30 min. Initial screening of crystals was set up in Crystalquick 96-well X plates (Greiner Bio-One) with a Cartesian Robot using the nanoliter sitting-drop vapor-diffusion method, with 100 nL of protein plus 100 nL of reservoir in each drop for Wuhan RBD/mAb-58/mAb-158 and Wuhan RBD/EY6A/beta-55 complexes, and 200 nL of protein plus 100 nL of reservoir for Omicron RBD/EY6A/beta-55 complex, as previously described (Walter et al., 2003). Crystals of Wuhan RBD/mAb-58/mAb-158 were formed in Hampton Research PEGRx condition 2-28, containing 0.1 M sodium citrate tribasic, pH 5.5 and 20% (w/v) PEG 4000. Crystals of Wuhan RBD/EY6A/beta-55 complex were obtained from Emerald Biostructures Wizard condition II- 7, containing 0.2 M NaCl, 0.1 M Tris, pH 6.7 and 30% (w/v) PEG 3000. Crystals of Omicron RBD/EY6A/beta-55 complex were formed in Hampton Research Index condition 80, containing 0.2 M (NH4)2COOH, 0.1 M Hepes, pH 7.5 and 25% (w/v) PEG 3350.

#### X-ray data collection, structure determination and refinement

Crystals were mounted in loops and dipped in solution containing 25% glycerol and 75% mother liquor for a second before being frozen in liquid nitrogen. Diffraction data were collected at 100 K at beamline I03 of Diamond Light Source, UK. All data were collected as part of an automated queue system allowing unattended automated data collection (https://www.diamond.ac.uk/Instruments/Mx/I03/I03-Manual/Unattended-Data-Collections.html). Diffraction images of 0.1° rotation were recorded on an Eiger2 XE 16M detector (exposure time from 0.02 to 0.03 s per image, beam size 80×20 μm or 50×20 μm, 10% to 30% beam transmission and wavelength of 0.9763 Å). Data were indexed, integrated and scaled with the automated data processing program Xia2-dials (Winter, 2010; Winter et al., 2018). 720° of data was collected from a crystal of Omicron-RBD/Beta-55/EY6A. 360° of data was collected for each of the Wuhan RBD/Beta-55/EY6A and Wuhan RDB/mAb-58/mAb- 158 data sets.

Structures were determined by molecular replacement with PHASER (McCoy et al., 2007). VhVl and ChCl domains which have the most sequence similarity to previously determined SARS-CoV-2 RBD/Fab structures (Dejnirattisai et al., 2021a; Dejnirattisai et al., 2021b; Huo et al., 2020; Liu et al., 2021a; Supasa et al., 2021; Zhou et al., 2021; Zhou et al., 2020) were used as search models for each of the current structure determination. Model rebuilding with COOT (Emsley et al., 2010) and refinement with Phenix (Liebschner et al., 2019) were used for all the structures. Data collection and structure refinement statistics are given in **Table S2**. Structural comparisons used SHP(Stuart et al., 1979), residues forming the RBD/Fab interface were identified with PISA (Krissinel and Henrick, 2007) and figures were prepared with PyMOL (The PyMOL Molecular Graphics System, Version 1.2r3pre, Schrödinger, LLC).

### QUANTIFICATION AND STATISTICAL ANALYSIS

Statistical analyses are reported in the results and figure legends. Neutralization was measured by FRNT. The percentage of focus reduction was calculated and IC50 (FRNT50) was determined using the probit program from the SPSS package. The Wilcoxon matched-pairs signed rank test was used for the analysis and two-tailed P values were calculated on geometric mean values.

Video S1. Antigenic map for major variants of SARS-CoV-2. Related to Figure 7.

