## Supplementary figures and tables for "Omicron-B.1.1.529 leads to widespread escape from neutralizing antibody responses"

A

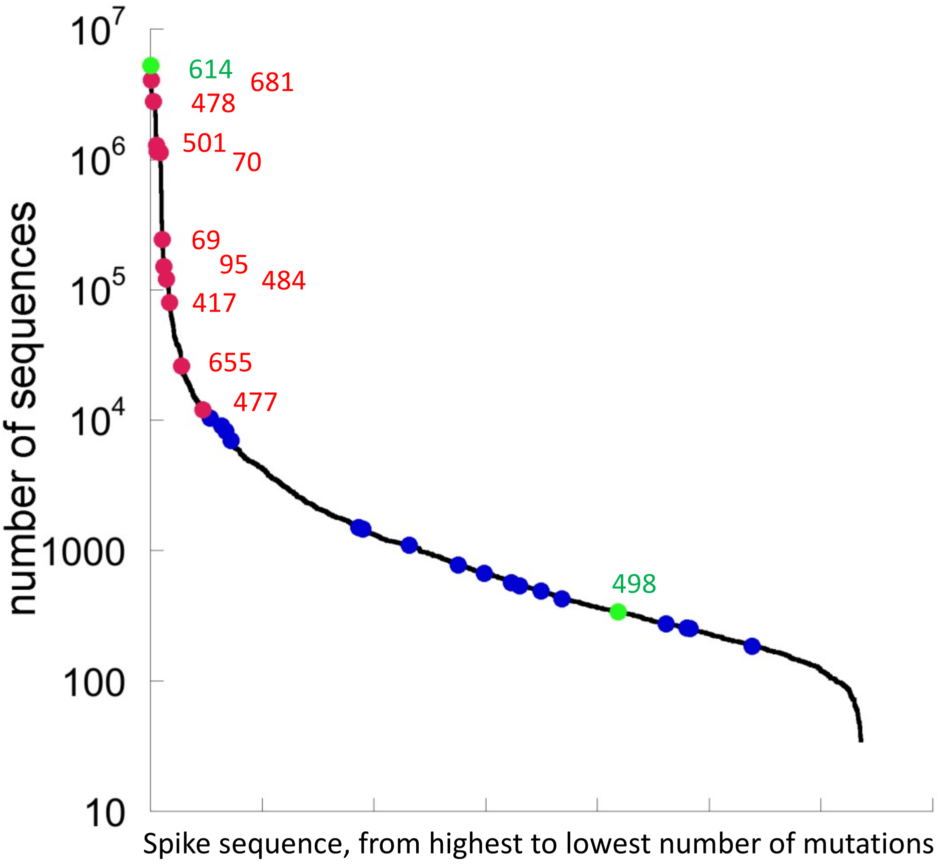


B

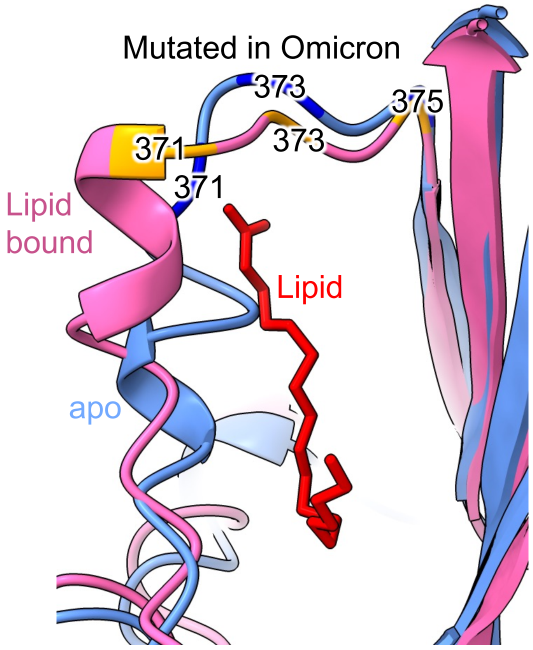


**Figure S1. A.** Number of sequenced mutations per position. The line shows the number of mutations per residue, for high to low along the spike protein. In green are mutations D614G, which is fixed from early virus evolution and position 498, which became dominant only in omicron. Red are for mutations in Omicron that were identified before in multiple linages and blue are mutations with Omicron being the only lineage.
**B.** Location of the S371L, S373P and S375F mutations in the context of the conformation change occurring on binding lipid. Cartoons of the apo (blue) and lipid bound (pink) early pandemic RBD are shown. The lipid is shown in red. Related to Figures 2 and 7E.

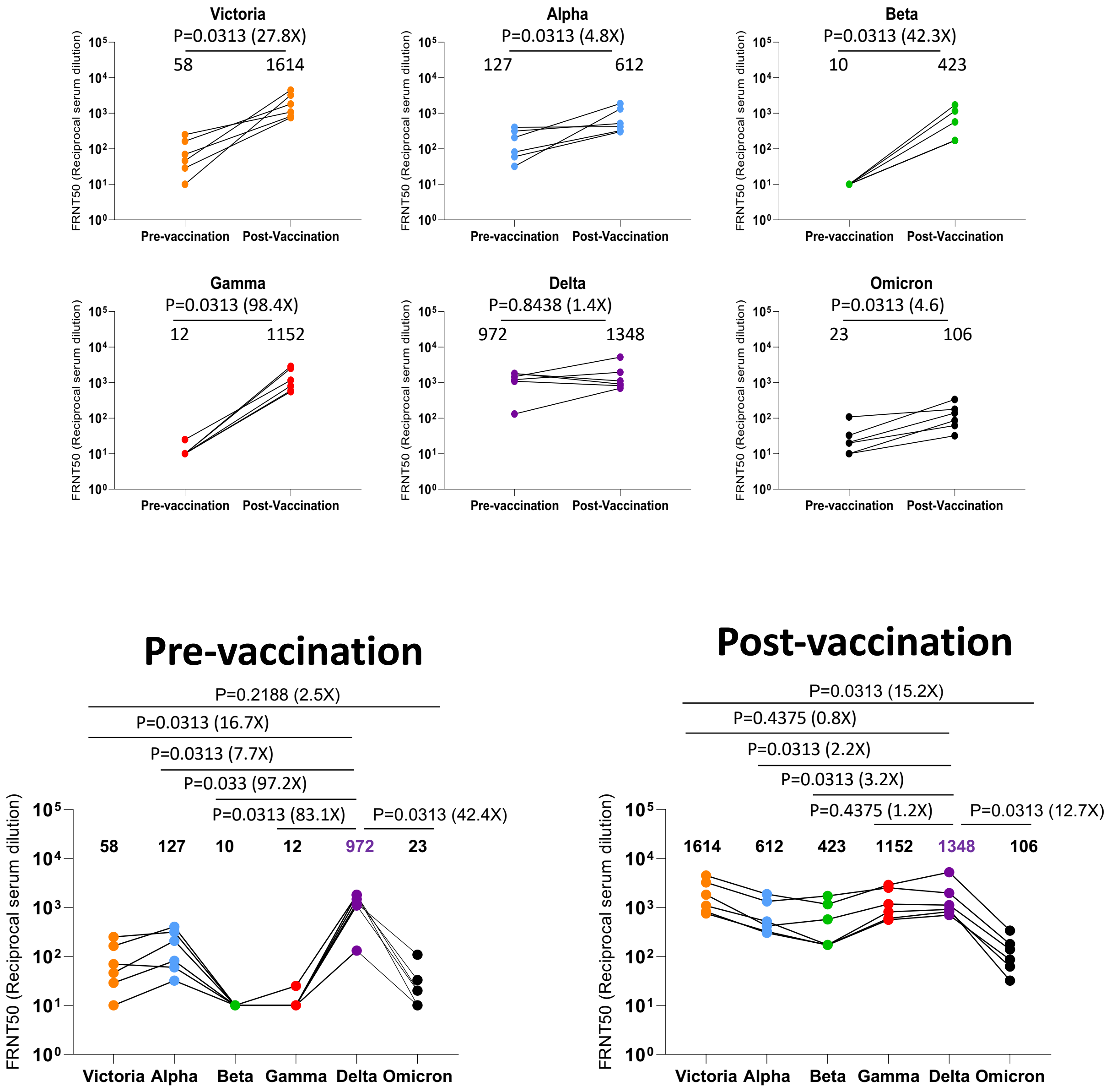


**Figure S2.** FRNT50 values for 7 cases of Delta infection before and after vaccination. Related to Figure 4.

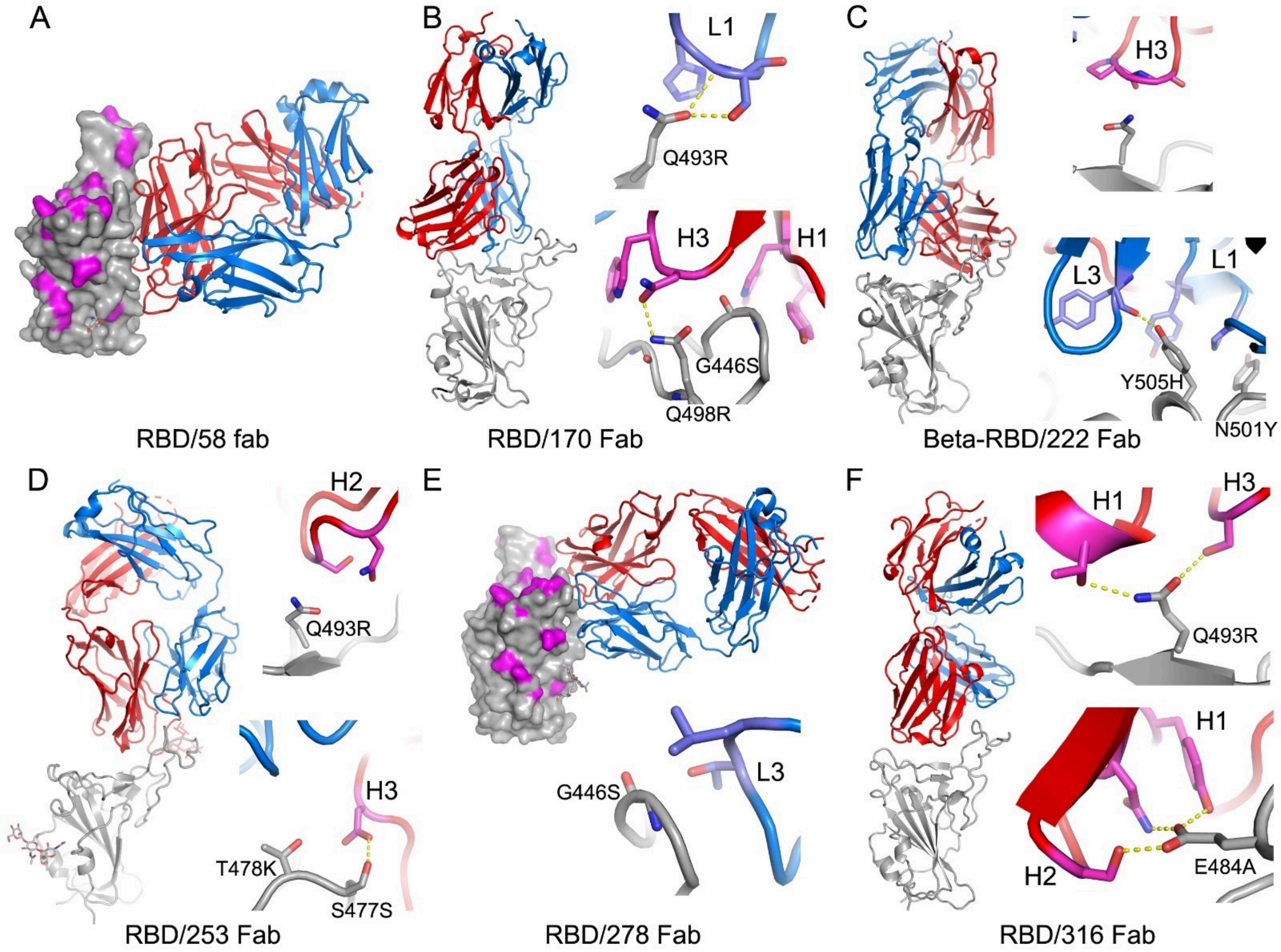


**Figure S3. Binding modes of early pandemic mAbs and their contacts to Omicron mutation sites.** Fabs are drawn as ribbons with the heavy chains in red and light chains in blue, and RBDs as grey ribbon or surface representation with Omicron mutation sites highlighted in magenta. Side chains are shown as sticks, and hydrogen bonds as dashed lines. (A) Fab 58 dosen't make any close contacts with the Omicron mutation sites. (B)-(F) Binding modes and contacts with Omicron mutation sites of Fabs 170, 222, 253, 278 and 316 respectively. Related to Figure 5.

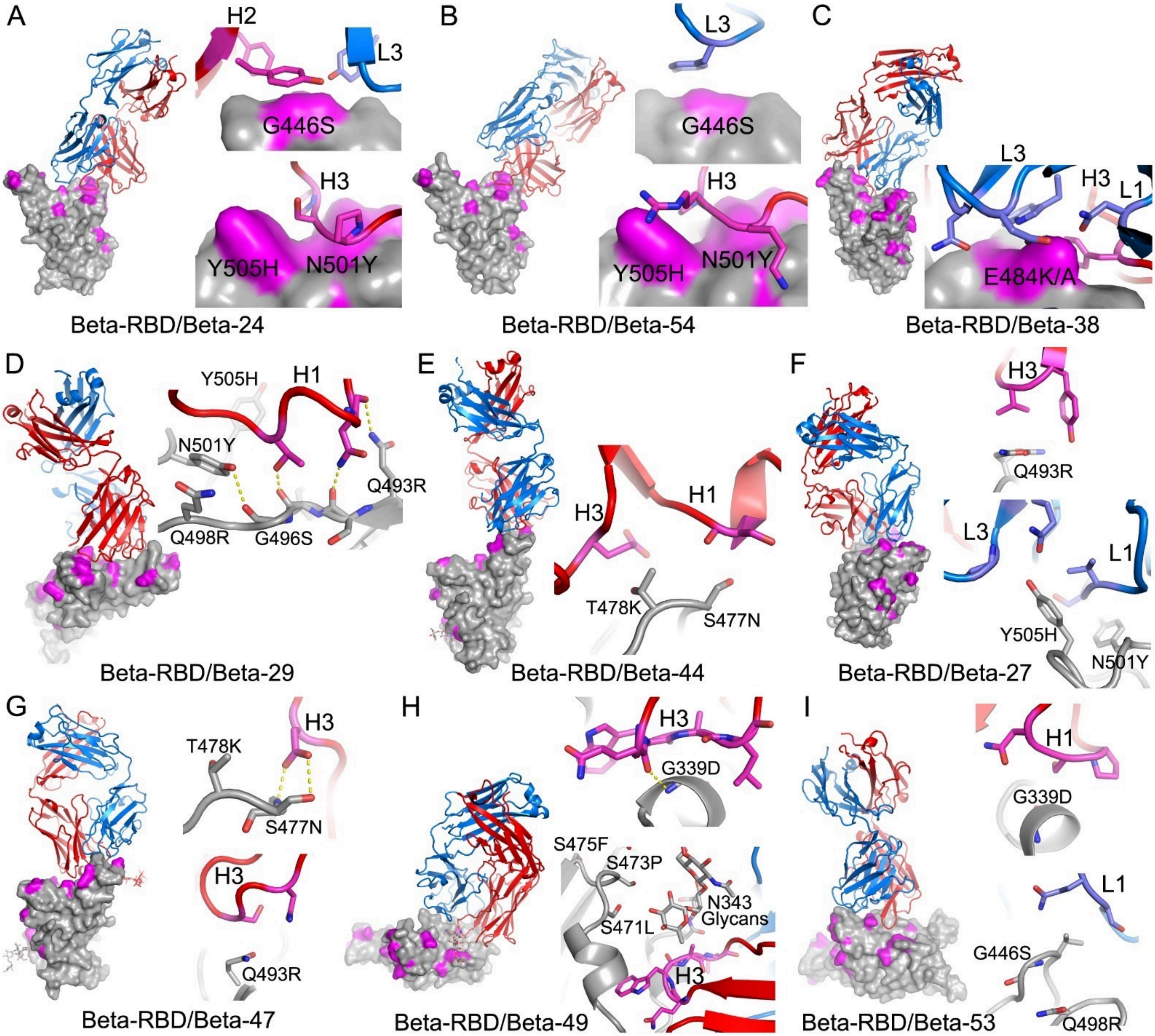


**Figure S4. Binding modes of Beta mAbs and their contacts to Omicron mutation sites.** The drawing and colouring schemes are same as in Figure S2. These are structures of Beta-RBD/Beta-Fab complexes. (A) Beta-24 and (B) Beta-54, examples of Beta mAbs targeting the N501Y mutation site. (C) Beta-38, a representative of Beta mAbs targeting the E484K mutation site. (D) Beta-29, a K417N/T dependent Beta mAb. (E) Beta-44 binds at the top of left shoulder and is sensitive to T478K mutation. (F)-(I) Beta-27, 47, 49 and 53 respectively. These four Beta mAbs neutralise all the previous variants of concern as well as the early pandemic Wuhan strain. Related to Figure 5.

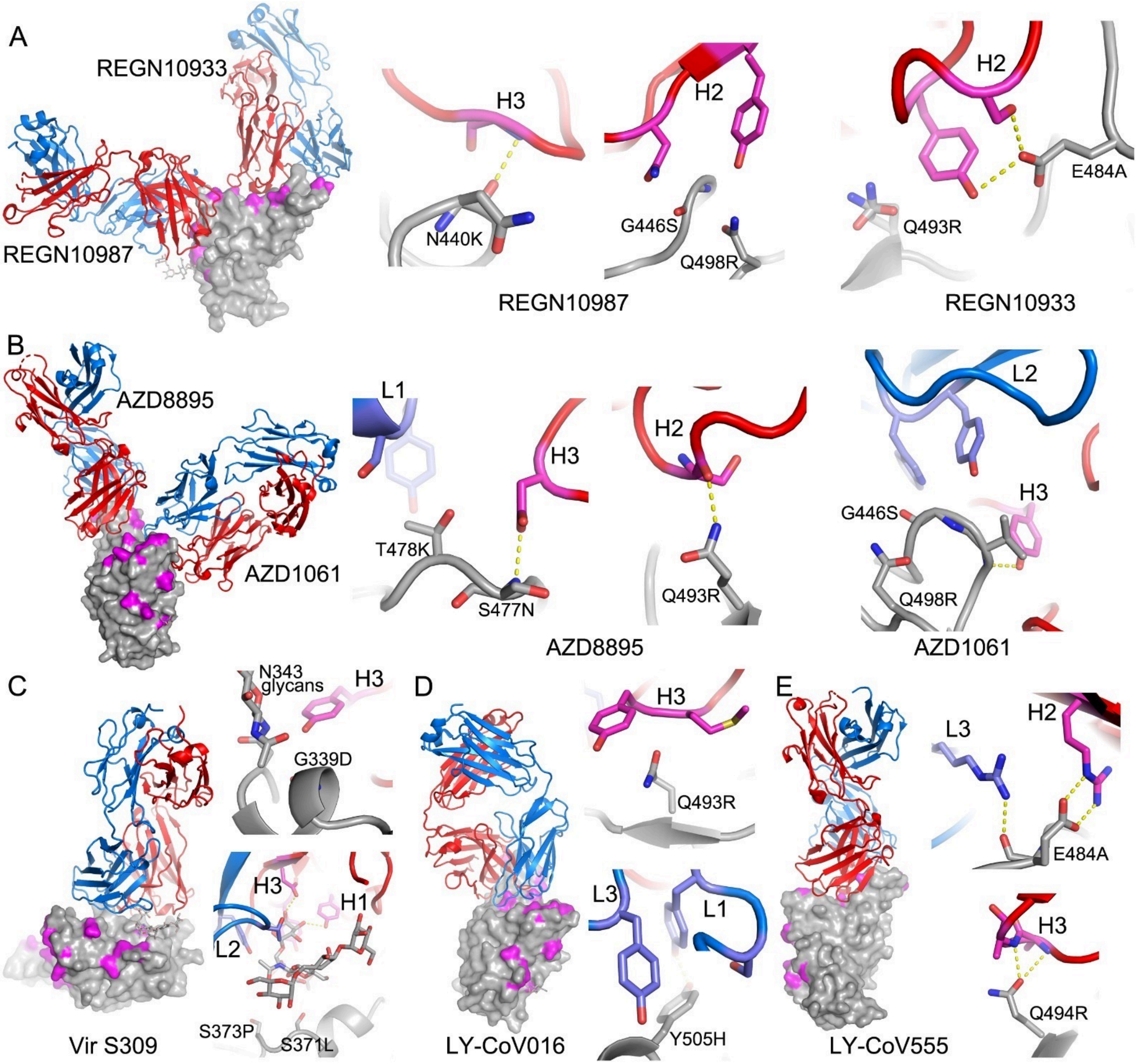


**Figure S5. Binding modes of the therapeutic mAbs and their contacts to Omicron mutation sites.** The drawing and colouring schemes are same as in Figure S2. (A) REGN10987 and REGN10933. (B) AZD8895 and AZD1061. (C) Vir S309. (D) LY-CoV016 and (E) LY-CoV555. Related to Figure 5.

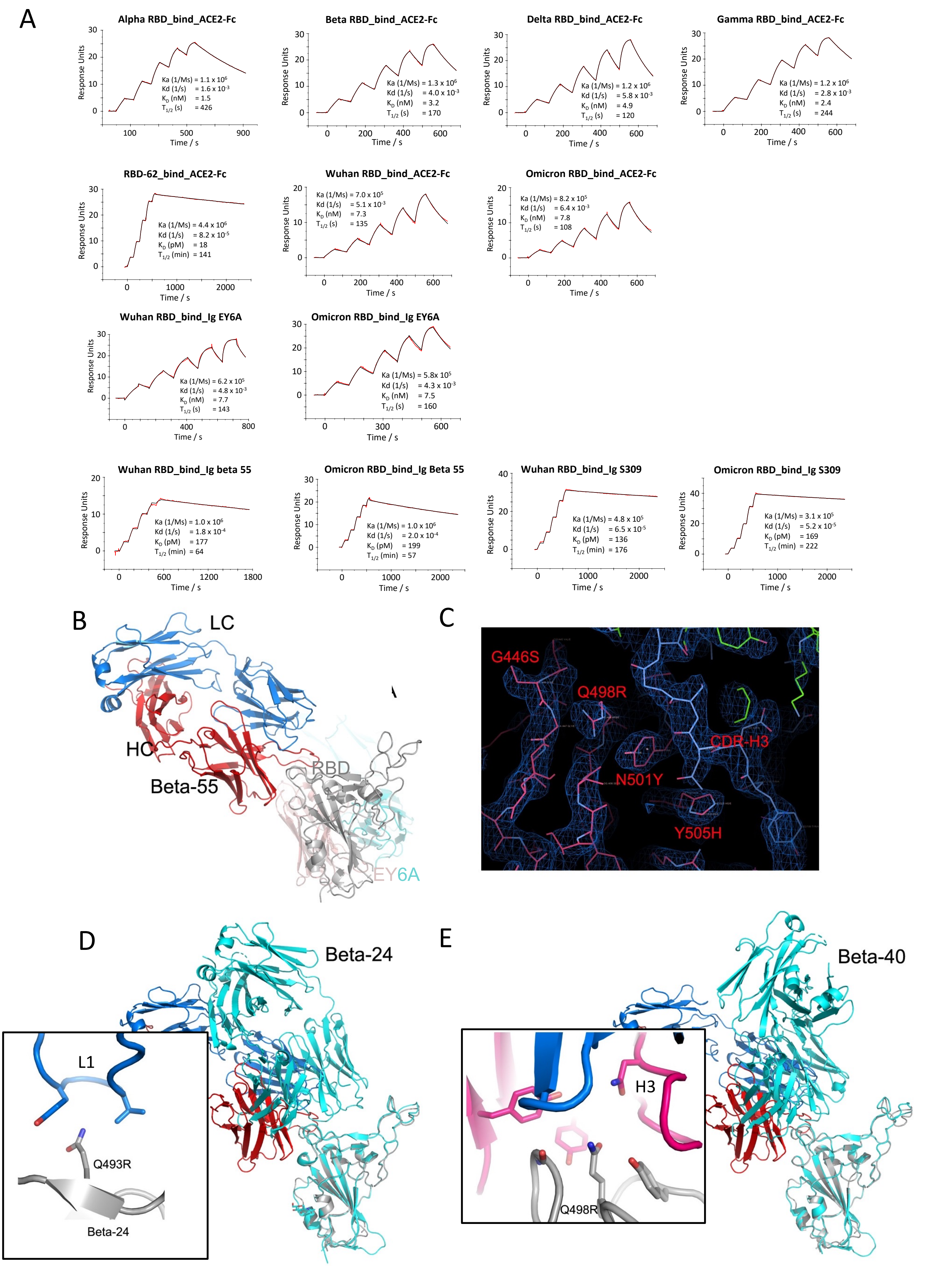


**Figure S6.** SPR measurement and Crystal structure of the Omicron RBD complexed with Beta-55 and EY6A Fabs. (A) SPR measuremments. (B) Ternary complex of the Omicron-RBD (grey)/Beta-55 (HC red, LC blue)/EY6A (HC salmon, LC cyan). (C) Electron density map showing the density for the mutated residues at 446, 498, 501 and 505, and their interactions with the CDR-H3 of Beta-55. (D) and (E) Comparison of the slightly different binding mode of Beta-55 to Beta-24 (cyan in (D)) and Beta-40 (cyan in (E)), the close-up boxes show details of the interactions with Beta-24 and Beta-40 explaining the knock-out of Beta-24 and the resilience of Beta-40. Related to Figure 7.

| mAb | IC50 (ug/ml) | | | | | |
|---|---|---|---|---|---|---|
| | Victoria | Alpha | Beta | Gamma | Delta | Omicron |
| 40 | 0.026 ± 0.007 | 0.035 ± 0.008 | 0.738 ± 0.311 | 0.153 ± 0.037 | 0.029 ± 0.010 | 7.989±2.011 |
| 55 | 0.095 ± 0.015 | 0.348 ± 0.044 | 0.127 ± 0.014 | 0.306 ± 0.046 | 0.016 ± 0.005 | 7.12±2.880 |
| 58 | 0.041 ± 0.003 | 0.116 ± 0.029 | 0.136 ± 0.010 | 0.236 ± 0.075 | 6.434 ± 2.623 | 0.141±0.063 |
| 88 | 0.033 ± 0.001 | 0.058 ± 0.008 | >10 | >10 | 0.039 ± 0.007 | >10 |
| 132 | 0.048 ± 0.000 | 0.337 ± 0.048 | >10 | >10 | 0.051 ± 0.013 | >10 |
| 150 | 0.012 ± 0.000 | 0.139 ± 0.019 | 0.350 ± 0.010 | 0.040 ± 0.003 | 0.020 ± 0.001 | >10 |
| 158 | 0.031 ± 0.004 | 0.254 ± 0.109 | >10 | >10 | 0.026 ± 0.002 | >10 |
| 159 | 0.011 ± 0.000 | 0.061 ± 0.020 | >10 | 1.434 ± 0.804 | >10 | >10 |
| 165 | 0.034 ± 0.004 | 0.212 ± 0.004 | 0.054 ± 0.013 | 0.241 ± 0.030 | 0.027 ± 0.006 | >10 |
| 170 | 0.025 ± 0.004 | 0.105 ± 0.050 | >10 | >10 | 0.841 ± 0.103 | >10 |
| 175 | 0.026 ± 0.000 | 0.575 ± 0.280 | >10 | 3.881 ± 0.738 | 0.017 ± 0.003 | >10 |
| 222 | 0.019 ± 0.000 | 0.014 ± 0.002 | 0.017 ± 0.005 | 0.008 ± 0.003 | 0.018 ± 0.001 | 0.240±0.122 |
| 253 | 0.055 ± 0.008 | 0.126 ± 0.018 | 0.109 ± 0.055 | 0.137 ± 0.005 | 0.005 ± 0.001 | 1.063±0.367 |
| 269 | 0.030 ± 0.000 | >10 | >10 | >10 | 0.021 ± 0.004 | >10 |
| 278 | 0.014 ± 0.007 | 0.307 ± 0.149 | 0.160 ± 0.018 | 0.245 ± 0.042 | 7.374 ± 1.397 | >10 |
| 281 | 0.005 ± 0.001 | 0.012 ± 0.000 | >10 | >10 | 1.494 ± 0.302 | >10 |
| 316 | 0.018 ± 0.007 | 0.024 ± 0.005 | >10 | >10 | 0.008 ± 0.001 | >10 |
| 318 | 0.029 ± 0.008 | 0.185 ± 0.037 | 0.019 ± 0.008 | 0.083 ± 0.032 | 0.018 ± 0.003 | >10 |
| 384 | 0.004 ± 0.001 | 0.005 ± 0.002 | >10 | >10 | 0.108 ± 0.035 | >10 |
| 398 | 0.091 ± 0.004 | 0.180 ± 0.001 | >10 | >10 | 0.237 ± 0.038 | >10 |
| 253-55 | 0.003 ± 0.000 | 0.008 ± 0.002 | 0.009 ± 0.002 | 0.026 ± 0.006 | 0.003 ± 0.000 | 2.945±1.283 |
| 253-165 | 0.003 ± 0.000 | 0.006 ± 0.000 | 0.013 ± 0.003 | 0.019 ± 0.000 | 0.007 ± 0.002 | >10 |
| β06 | >10 | 0.024 ± 0.002 | 0.008 ± 0.002 | 0.015 ± 0.003 | >10 | >10 |
| β10 | >10 | 0.064 ± 0.042 | 0.015 ± 0.000 | 0.025 ± 0.011 | >10 | >10 |
| β20 | >10 | >10 | 0.005 ± 0.001 | 0.345 ± 0.122 | >10 | 7.518±1.105 |
| β22 | >10 | 6.58 ± 2.988 | 0.025 ± 0.004 | 0.030 ± 0.007 | >10 | 0.393±0.234 |
| β23 | >10 | 0.009 ± 0.001 | 0.011 ± 0.001 | 0.020± 0.000 | >10 | >10 |
| β24 | >10 | 0.007 ± 0.001 | 0.002 ± 0.001 | 0.005 ± 0.001 | >10 | >10 |
| β26 | 2.742 ± 0.208 | >10 | 0.012 ± 0.003 | 0.016 ± 0.000 | >10 | >10 |
| β27 | 0.018 ± 0.002 | 0.018 ± 0.000 | 0.009 ± 0.000 | 0.006 ± 0.002 | 0.021 ± 0.004 | 2.693±0.741 |
| β29 | >10 | 1.372 ± 0.016 | 0.027 ± 0.003 | 0.023 ± 0.009 | >10 | 0.261±0.079 |
| β30 | 2.643 ± 0.88 | 0.004 ± 0.001 | 0.003 ± 0.001 | 0.004 ± 0.000 | 0.350 ± 0.035 | >10 |
| β32 | 0.248 ± 0.003 | 0.119 ± 0.044 | 0.053 ± 0.025 | 0.027 ± 0.014 | 0.267 ± 0.068 | >10 |
| β33 | 2.016 ± 0.051 | 0.234 ± 0.013 | 0.017 ± 0.003 | 0.017 ± 0.001 | 0.334 ± 0.005 | >10 |
| β34 | 8.241 ± 1.067 | 1.466 ± 0.136 | 0.032 ± 0.010 | 0.092 ± 0.003 | >10 | >10 |
| β38 | >10 | >10 | 0.011 ± 0.003 | 0.043 ± 0.025 | >10 | >10 |
| β40 | 0.075 ± 0.005 | 0.001 ± 0.000 | 0.001 ± 0.000 | 0.001 ± 0.000 | 0.107 ± 0.031 | 0.012±0.007 |
| β43 | >10 | >10 | 0.048±0.024 | >10 | >10 | >10 |
| β44 | 0.007 ± 0.002 | 0.028 ± 0.008 | 0.015 ± 0.008 | 0.071 ± 0.026 | >10 | >10 |
| β45 | >10 | >10 | 0.018 ± 0.003 | 0.015 ± 0.006 | >10 | 9.947±0.053 |
| β47 | 0.006 ± 0.001 | 0.008 ± 0.003 | 0.003 ± 0.001 | 0.004 ± 0.002 | 0.005 ± 0.000 | 0.096±0.011 |
| β48 | 0.034 ± 0.011 | 0.020 ± 0.008 | 0.009 ± 0.001 | 0.011 ± 0.001 | 0.042 ± 0.016 | 8.362±1.638 |
| β49 | 0.009 ± 0.000 | 0.011 ± 0.001 | 0.007 ± 0.000 | 0.019 ± 0.003 | 0.008 ± 0.003 | >10 |
| β50 | 0.011 ± 0.000 | 0.014 ± 0.006 | 0.007 ± 0.001 | 0.015 ± 0.005 | 0.019 ± 0.005 | >10 |
| β51 | 0.119 ± 0.008 | 0.242 ± 0.024 | 0.005 ± 0.000 | 0.019 ± 0.001 | >10 | >10 |
| β53 | 0.005 ± 0.000 | 0.032 ± 0.009 | 0.004 ± 0.000 | 0.017 ± 0.001 | 0.007 ± 0.000 | 0.266±0.117 |
| β54 | 0.232 ± 0.092 | 0.002 ± 0.001 | 0.001 ± 0.000 | 0.002 ± 0.001 | 0.409 ± 0.071 | 0.020±0.011 |
| β55 | 0.108 ± 0.069 | 0.028 ± 0.004 | 0.01 0± 0.003 | 0.022 ± 0.001 | 0.076 ± 0.020 | 0.109±0.013 |
| β56 | 0.046 ± 0.013 | 0.001 ± 0.000 | 0.001 ± 0.000 | 0.002 ± 0.000 | 0.022 ± 0.002 | 0.194±0.056 |
| AZD1061 | 0.013 ± 0.003 | 0.012 ± 0.002 | 0.014 ± 0.002 | 0.007 ± 0.002 | 0.038 ± 0.006 | 3.488±2.085 |
| AZD8895 | 0.005 ± 0.001 | 0.011 ± 0.002 | 0.046 ± 0.031 | 0.046 ± 0.016 | 0.003 ± 0.000 | 1.152±0.170 |
| AZD7442 | 0.009 ± 0.000 | 0.007 ± 0.001 | 0.012 ± 0.001 | 0.006 ± 0.003 | 0.005 ± 0.000 | 0.273±0.062 |
| REGN10987 | 0.032 ± 0.007 | 0.028 ± 0.003 | 0.007 ± 0.001 | 0.013 ± 0.002 | 0.017 ± 0.009 | >10 |
| REGN10933 | 0.004 ± 0.002 | 0.014 ± 0.002 | 3.284 ± 2.014 | 6.177 ± 1.914 | 0.003 ± 0.001 | >10 |
| ADG10 | 0.006 ± 0.000 | 0.010 ± 0.001 | 0.011 ± 0.001 | 0.003 ± 0.000 | 0.026 ± 0.005 | >10 |
| ADG20 | 0.004 ± 0.001 | 0.006 ± 0.000 | 0.01 ± 0.001 | 0.009 ± 0.000 | 0.006 ± 0.001 | 1.104±0.509 |
| ADG30 | 0.007 ± 0.002 | 0.016 ± 0.001 | 0.029 ± 0.003 | 0.002 ± 0.001 | 0.033 ± 0.007 | >10 |
| Ly-CoV-555 | 0.006 ± 0.002 | 0.009 ± 0.000 | >10 | >10 | 8.311 ± 4.059 | >10 |
| Ly-CoV16 | 0.034 ± 0.007 | 3.225 ± 1.030 | >10 | >10 | 0.012 ± 0.002 | >10 |
| S309 | 0.040 ± 0.005 | 0.078 ± 0.069 | 0.082 ± 0.002 | 0.076 ± 0.014 | 0.113 ± 0.028 | 0.256±0.034 |

**Table S1.** FRNT50 data, related to Figure 4.

| **Structure** | RBD/58-158 | RBD/Beta-55-EY6A | Omicron-RBD/ Beta-55-EY6A |
|---|---|---|---|
| **Data collection** | | | |
| Space group | $P3_121$ | $P3_221$ | $P3_221$ |
| Cell dimensions | | | |
| *a*, *b*, *c* (Å) | 72.8, 72.8, 416.2 | 131.8, 131.8, 116.8 | 132.0, 132.0, 117.3 |
| a, b, g (°) | 90, 90, 120 | 90, 90, 120 | 90, 90, 120 |
| Resolution (Å) | 104–2.84 (2.89–2.84) | 82–2.92 (2.97–2.92) | 117–2.40 (2.44–2.40) |
| $R_{merge}$ | 0.290 (---) | 0.554 (---) | 0.339 (---) |
| $R_{pim}$ | 0.071 (0.737) | 0.125 (0.902) | 0.053 (0.458) |
| $I/s(I)$ | 5.0 (0.3) | 3.8 (0.5) | 8.5 (0.7) |
| $CC_{1/2}$ | 0.996 (0.285) | 0.986 (0.311) | 0.977 (0.319) |
| Completeness (%) | 100 (100) | 99.9 (96.5) | 100 (100) |
| Redundancy | 17.9 (19.3) | 20.5 (20.2) | 41.1 (42.0) |
| **Refinement** | | | |
| Resolution (Å) | 70–2.84 | 82–2.92 | 114–2.40 |
| No. reflections | 29939/1578 | 24493/1236 | 44201/2278 |
| $R_{work}$ / $R_{free}$ | 0.229/0.279 | 0.223/0.268 | 0.210/0.253 |
| No. atoms | | | |
| Protein | 7933 | 8097 | 8126 |
| Ligand/ion/water | 36 | 28 | 144 |
| *B* factors (Å$^2$) | | | |
| Protein | 116 | 74 | 65 |
| Ligand/ion/water | 114 | 119 | 70 |
| r.m.s. deviations | | | |
| Bond lengths (Å) | 0.002 | 0.002 | 0.002 |
| Bond angles (°) | 0.5 | 0.5 | 0.5 |

**Table S2**. X-ray data collection and structure refinement statistics. Related to Figure 7.
